# The Mitochondrial RNA Granule Modulates Manganese-Dependent Cell Toxicity

**DOI:** 10.1101/2022.01.04.474973

**Authors:** E. Werner, A. Gokhale, M. Ackert, C. Xu, Z. Wen, A. M. Roberts, B. R. Roberts, A. Vrailas-Mortimer, A. Crocker, V. Faundez

## Abstract

Prolonged manganese exposure causes manganism, a neurodegenerative movement disorder. The identity of adaptive and non-adaptive cellular processes targeted by manganese remains mostly unexplored. Here we study mechanisms engaged by manganese in genetic cellular models known to increase susceptibility to manganese exposure, the plasma membrane manganese efflux transporter SLC30A10 and the mitochondrial Parkinson’s gene PARK2. We found that SLC30A10 and PARK2 mutations as well as manganese exposure compromised the mitochondrial RNA granule as well as mitochondrial transcript processing. These RNA granule defects led to impaired assembly and function of the mitochondrial respiratory chain. Notably, cells that survived a cytotoxic manganese challenge had impaired RNA granule function, thus suggesting that this granule phenotype was adaptive. CRISPR gene editing of subunits of the mitochondrial RNA granule, FASTKD2 or DHX30, as well as pharmacological inhibition of mitochondrial transcription-translation, were protective rather than deleterious for survival of cells acutely exposed to manganese. Similarly, adult *Drosophila* mutants with defects in the mitochondrial RNA granule component *scully* were safeguarded from manganese-induced mortality. We conclude that the downregulation of the mitochondrial RNA granule function is a protective mechanism for acute metal toxicity.

**Significance Statement:** Mutations in the manganese efflux transporter SLC30A10 and the mitochondrial Parkinson’s gene PARK2, cause neurodegeneration and increased susceptibility to toxic manganese exposure. Thus, molecular processes affected in both mutants could offer insight into fundamental mechanisms conferring susceptibility or resilience to environmental and genetic factors associated with neurodegeneration. Here we report that SLC30A10 and PARK2 mutations compromise an understudied structure, the mitochondrial RNA granule, which is required for processing polycistronic mitochondrial RNAs. Cells and *Drosophila* lacking mitochondrial RNA granule components were resistant to manganese exposure. We conclude that the downregulation of the mitochondrial RNA granule function is an adaptive mechanism for cells exposed to manganese.

## Introduction

Manganese accumulation in the brain results in manganism, a parkinsonian neurodegenerative disorder (1–5). Manganism and Parkinson’s disease are distinguishable clinically and considered distinct movement disorders. However, epidemiological and *in vitro* evidence suggest a degree of overlap between these two neurodegenerative diseases (2, 5). Human environmental exposure studies suggest that manganese could increase the risk of Parkinson’s disease (6–9). Moreover, genetic defects in Parkinson’s disease causative genes increase cellular manganese susceptibility and manganese exposure modifies the function of products encoded by the Parkinson’s disease genes PARK1 and PARK2 (10–13). Nevertheless, evidence for convergent mechanisms common to manganism and Parkinson’s disease remains elusive. Here, we take comprehensive, unbiased, molecular systems approaches to identify pathways affected in mutant cellular models, which are known to increase manganese susceptibility. We focused on the SLC30A10 and PARK2 genes whose mutations increase manganese susceptibility while causing a manganism-type syndrome and Parkinson’s disease, respectively.

SLC30A10 is a plasma-membrane localized efflux transporter necessary for cellular manganese homeostasis (14, 15). Genetic defects in SLC30A10 results in hypermanganesemia, dystonia, and parkinsonism (OMIM: 613280), offering an autosomal recessive inheritable manganese toxicity model(15–18). PARK2 encodes a mitochondrially localized ubiquitin E3 ligase required for mitophagy (19–21). PARK2 mutations cause autosomal recessive juvenile Parkinson’s disease (OMIM: 613280) (22–24). Importantly, some effects of manganese exposure on mitochondria mimic what has been reported in genetic forms of Parkinson’s disease (25). Moreover, mutations in PARK2 increase manganese susceptibility {Chakraborty, 2015 #297;Bornhorst, 2014 #298}. Thus, although SLC30A10 and PARK2 reside in separate compartments, we ask if there are shared molecular alterations between mutants of these two genes. We hypothesized that such novel and shared mechanisms in SLC30A10 and PARK2 mutant cell models could influence cellular fitness after metal exposure.

We report our comparative molecular systems studies in SLC30A10 and PARK2 mutant cells, which identified FASTKD2 as a factor whose expression was compromised in both mutant genotypes. FASTKD2 is a component of the mitochondrial RNA granule required for the processing of polycistronic mitochondrial RNAs, a step necessary for mitochondrial protein synthesis (26–28). We demonstrate that SLC30A10 and PARK2 mutations compromise mitochondrial RNA processing and, consequently, the assembly and function of respiratory chain complexes. Remarkably, genetic disruption of the mitochondrial RNA granule or pharmacological inhibition of mitochondrial transcription-translation in cells were protective against acute manganese exposure. A similar protective effect was obtained in *Drosophila* mutants of *scully*, a component of the mitochondrial RNA granule was disrupted. Our findings are unprecedented as they identify the mitochondrial RNA granule and RNA processing as key protective mechanisms in acute manganese toxicity. Our results upend the conception that compromised mitochondrial function is pathogenic in manganese toxicity.

## Results

### Genome-Wide Expression Analysis of Manganese Toxicity

We performed genome wide gene expression analyses seeking mechanistic overlaps between gene defects in the manganese efflux transporter SLC30A10, causative of Parkinsonism (OMIM: 613280), and the Parkinson’s gene PARK2 (OMIM: 600116). We CRISPR edited either one of these genes in the human cell line HAP1. This is a near-haploid cell line with a doubling time of ~15 h, which makes it uniquely suitable for rapid turnover of mitochondria and the selection of toxicant-resistant cell populations(29). SLC30A10 and PARK2 null cells (*SLC30A10^Δ^* and *PARK2^Δ^* cells, respectively) have enhanced sensitivity to increasing concentrations of manganese as compared to wild type cells. Both null genotypes significantly interacted with manganese exposure (Fig. 1A, p<0.001 for *SLC30A10^Δ^* cells and p<0.011 for *PARK2^Δ^* cells, 2-Way ANOVA), as has been reported before (30). In contrast, exposure to copper did not reveal interactions between cell genotype and copper exposure. Despite similarities in manganese susceptibility, SLC30A10 and PARK2 null cells differed in their capacity of handling a manganese challenge. *SLC30A10^Δ^* cells significantly increased metal content after a manganese challenge while PARK2 mutants behaved as their wild type controls before or after manganese exposure (Fig. 1B).

**Figure 1.**
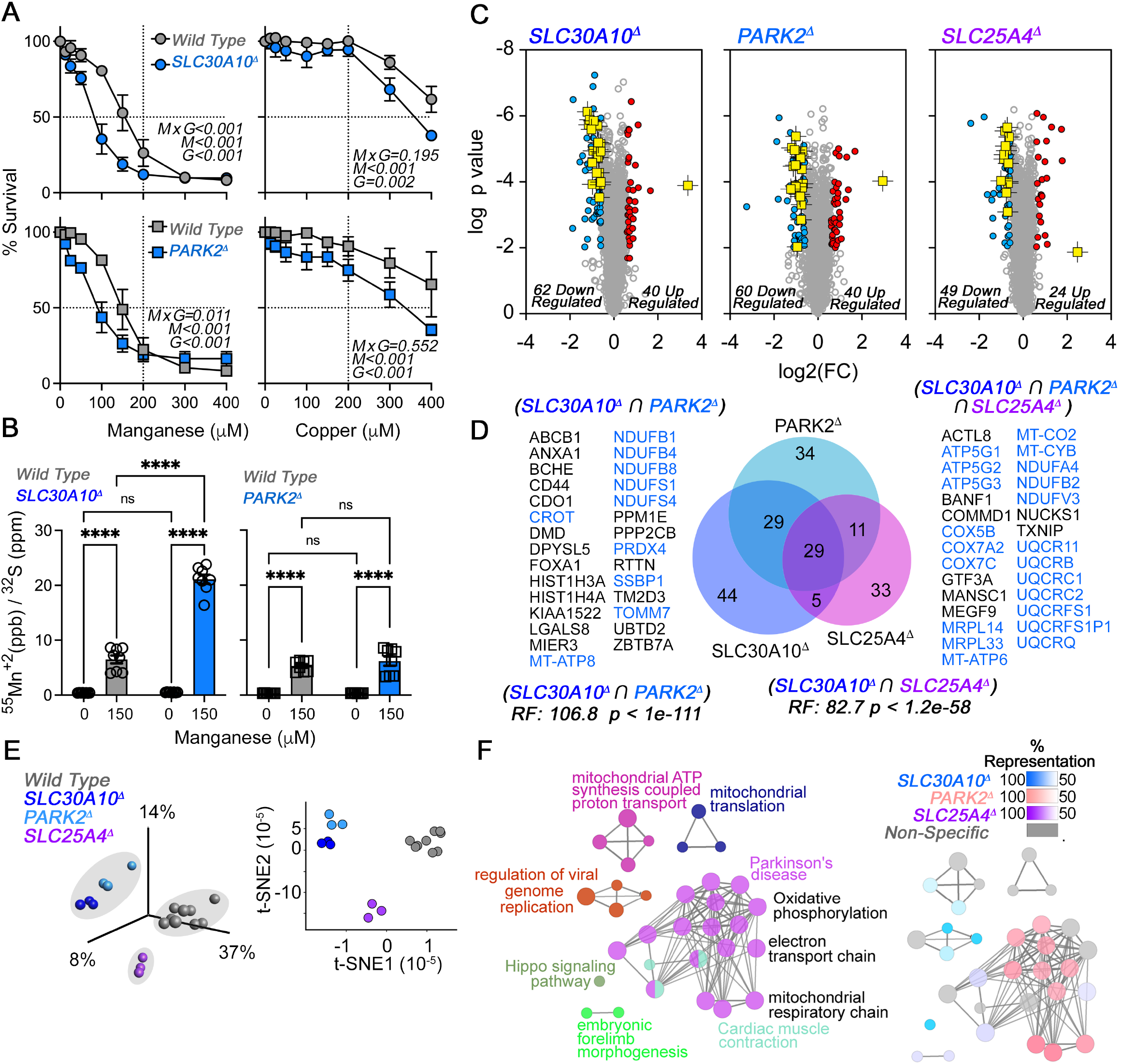
Comparative Proteomics Analysis of SLC30A10 and Mitochondrial Mutants. **A**. Cell survival analysis of SLC30A10 and PARK2 mutants with increasing concentrations of manganese or copper. Average ± SEM, n=3, Two-Way ANOVA followed by Benjamini, Krieger and Yekutieli corrections. **B**. ICP mass spectrometry determination of manganese in HAP1 cells treated with vehicle or manganese overnight. Data were normalized to total sulphur content. Average ± SEM, n=8, Two-Way ANOVA followed by Šydák’s corrections. **C**. Volcano plots of TMT labeled wild type HAP1 cells or mutants in either SLC30A10, PARK2, or SLC25A4. Red symbols denote upregulated proteins, blue symbols denote downregulated proteins and yellow symbols mark mitochondrial proteins significantly affected. n=3, depicted are Benjamini-Hochberg corrected log_10_ q values and log_2_ fold of change. **D**. Venn diagram of significant hits shared by the three mutant genotypes in C. All mitochondrial protein hits annotated in Mitocarta 3.0 are labeled in blue font. p value was calculated with exact hypergeometric probability. **E**. SLC25A4 mutant cells diverge from SLC30A10 and PARK2 null cells using Principal Component Analysis and 2D-tSNE. Close grouping of SLC30A10 and PARK2 mutants was determined by k-means clustering. **F**. Gene ontology analysis of all proteome hits in SLC30A10, PARK2, and SLC25A4 mutants. Overlapping and mutant-specific ontologies are color-coded by percent of contribution >50% to an ontological category. Gray represents ontologies where all three mutants similarly contribute hits.

We performed multiplexed quantitative Tandem Mass Tagging mass spectrometry to quantify the proteome in these SLC30A10 and PARK2 null cells. We wanted to unbiasedly assess proteome wide similarities/differences between these genotypes that could inform mechanistic and phenotypic convergence/divergence. We contrasted SLC30A10 and PARK2 null cell proteomes with an isogenic SLC25A4 mutant HAP1 cell line, affecting the mitochondrial ADP-ATP translocator. Our goal was to distinguish to what extent SLC30A10 and PARK2 mutant proteome similarities were merely a consequence of disrupted bioenergetics, which we modeled with the SLC25A4 mutation (Fig. 1C) (31). We simultaneously quantified 7,079 proteins in these three mutant genotypes, which correspond to 72% of the known proteome of the HAP1 parental cell line (32) (Supplementary Table 1). We identified between ~70-100 proteins whose content was modified by genotype. These three mutant proteomes converged on 29 proteins, 21 of which were annotated to mitochondria and of reduced expression (Fig. 1D, blue text). The overlap between the SLC30A10 and PARK2 mutant proteomes was higher than between SLC30A10 and SLC25A4 proteomes as quantified by the number of common hits and the representation factor, a figure that quantifies the extent of overlap between gene sets above what it is expected by chance (Fig. 1D, RF). Unique among the proteins whose expression was changed solely by SLC30A10 and PARK2 mutants was the upregulation of SSBP1 (Fig. 1C-D), a mitochondrial singlestranded DNA binding protein, required to maintain the mitochondrial genome and to process mitochondrial RNAs (33, 34). We further scrutinized the convergence of proteomes among mutant genotypes. Principal component analysis combined with k-mean clustering revealed that the proteomes of SLC30A10 and PARK2 mutant cells were closer to each other than to SLC25A4 mutant cells (Fig.1 E). A similar outcome was obtained with non-linear data dimensionality reduction with 2D-tSNE (Fig.1 E). Finally, we used gene ontology to search for biological processes preferentially affected by each of these three mutant genes. We found that the only specific ontologies were associated to *SLC30A10^Δ^* proteome (Supplementary Table 2). These ontologies correspond to a downregulation of the Hippo signaling pathway (Fig. 1F, KEGG:04392, p=0.0026, group p value Bonferroni corrected) and increase in the levels of proteins annotated to negative regulation of viral genome replication, the later represented by Interferon Induced Transmembrane Proteins IFITM1-3 (Fig. 1F, GO:0045071, p=0.00073, group p value Bonferroni corrected). Both of these pathways are required for a response to DNA and RNA viruses (35). Despite the divergence of the whole proteome in SLC30A10 and PARK2 mutants with the SLC25A4 deficient cells (Fig. 1E), we found that mitochondrial ontologies were similarly shared by the three gene defects studied (Fig. 1F). These results demonstrate that the SLC30A10 and PARK2 proteomes are closely related and distinct from the proteome of SLC25A4 mutant cells, a model of impaired mitochondrial bioenergetics.

Proteome overlap between SLC30A10 and PARK2 null cells could result from parallel changes in the transcriptome or proteome convergence could originate from affected transcripts not captured by our proteomes. Thus, we performed RNAseq to identify transcriptomes altered by SLC30A10 and PARK2 mutants (Fig. 2A and Supplementary table 3). There was limited transcript overlap as evidenced by PCI and 2D-tSNE analysis, which revealed that the *PARK2^Δ^* transcriptome was closer to its wild type controls than to the transcriptome of *SLC30A10^Δ^* cells (Fig. 2A-B). Even though there was an overlap of 120 transcripts between the mutant SLC30A10 and PARK2 transcriptomes, the magnitude of this overlap was 17 times lower than the overlap of the mutant proteomes (Fig. 2C, RF 6.2 compare with RF 106.8 in Fig. 1D). Moreover, and in contrast to the proteome of SLC30A10 and PARK2 mutant cells, mutant transcriptomes did not enrich mitochondrially annotated transcripts above what is expected by chance (Fig. 2C *SLC30A10^Δ^* ⋂Mitocarta, RF: 0.8 and *PARK2^Δ^* ⋂Mitocarta, RF: 0.5). The limited mitochondrial annotated transcripts found had no overlap with the mutant mitochondrial proteome, with one exception APEX2, whose protein and transcript levels were increased in *SLC30A10^Δ^* cells (Fig. 2C). The only common mitochondrial annotated transcript shared by *SLC30A10^Δ^* and *PARK2^Δ^* cells was FASTKD2 (Fig. 2A and C). FASTKD2 is required for mitochondrial RNA processing and its deficiency impacts mitochondrial protein synthesis resulting in an impaired electron transport chain (36, 37). FASTKD2 was downregulated in both *SLC30A10^Δ^* and *PARK2^Δ^* cells and was the most downregulated gene expressed in *SLC30A10^Δ^*, a phenotype we confirmed by qRT-PCR (Fig. 2A and D). FASTKD2 mRNA expression was four orders of magnitude lower in *SLC30A10^Δ^* cells as compared to wild type cells (Fig. 2D).

**Figure 2.**
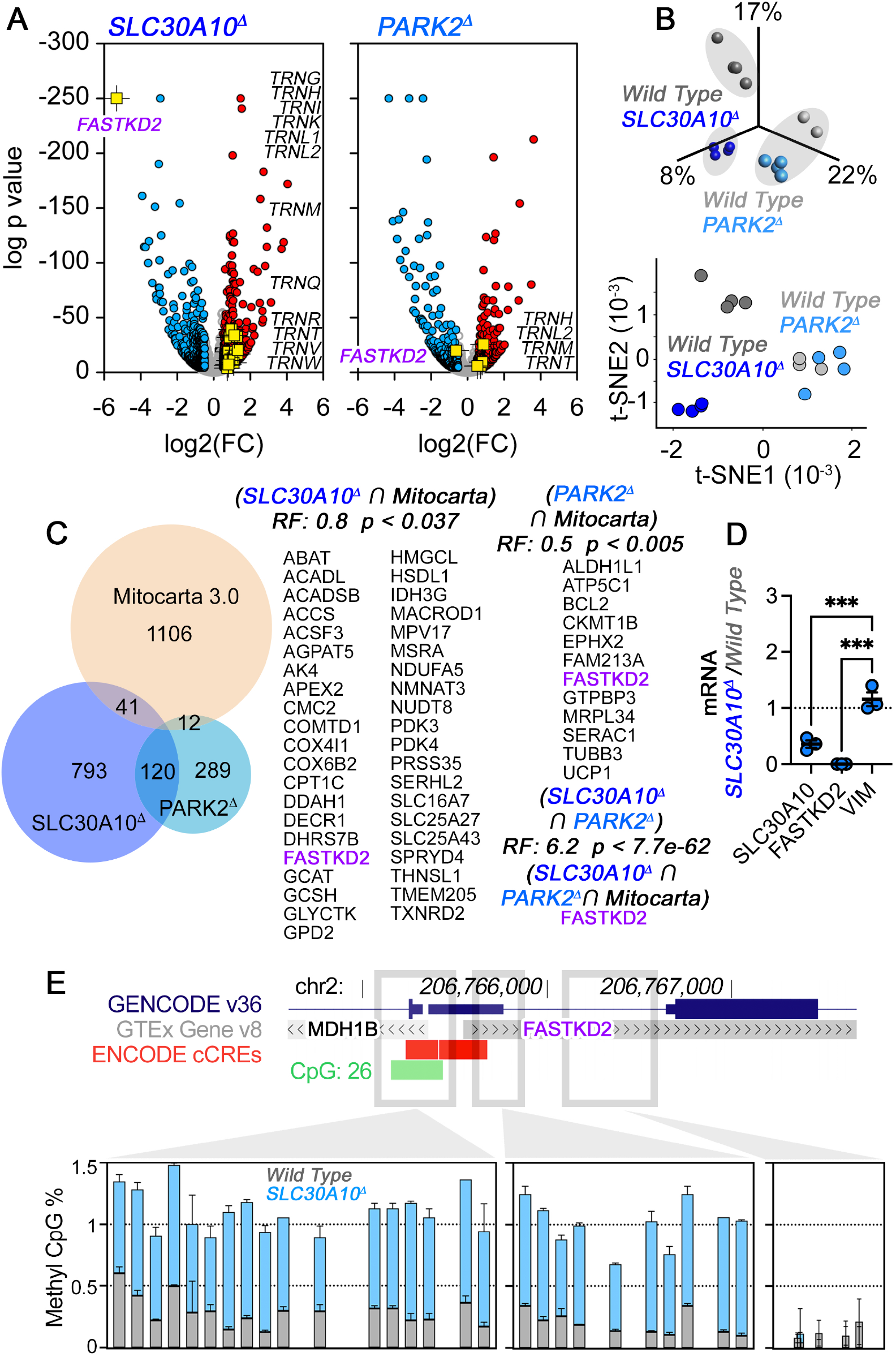
FASTKD2 and Mitochondrial tRNAs Belong to the Shared Transcriptome of SLC30A10 and PARK2 Mutants. **A**. Volcano plots of RNAseq experiments in wild type HAP1 cells, SLC30A10, or PARK2 mutants. Red symbols denote upregulated mRNAs, blue symbols denote downregulated mRNAs, and yellow symbols mark FASTKD2 and mitochondrial tRNAs listed in the plot. n=4, depicted are log_10_ p value calculated with Wald test corrected by Benjamini and Hochberg and log_2_ fold of change. **B**. Divergence of the SLC30A10 and PARK2 null cell transcriptome. Principal Component Analysis and 2D-tSNE analyses. Genotype grouping was determined by k-means clustering. **C**. Venn diagram of significant hits shared by SLC30A10 and PARK2 mutants annotated in Mitocarta 3.0. Only one Mitocarta 3.0 hit, FASTKD2, is common to SLC30A10 and PARK2 mutants. p value was calculated with exact hypergeometric probability. **D**. qRT-PCR analysis of SLC30A10 and FASTKD2 mRNAs in SLC30A10 mutant and wild type cells. Vimentin mRNA was used as control (VIM). One-Way ANOVA followed by Šydák’s corrections **E**. Methylation analysis of the FASTKD2 promoter region. UCSC Genome Map of the human FASTKD2 promoter region with annotated transcript (GTEx v8, gray), promoter-like signature (ENCODE, red), and CpG island (green). Graphs shows the percent methylation across amplicons for each genotype analyzed by Epityper. Average ± SEM, n=3.

We reasoned that the near-haploid genome organization in HAP1 cells could facilitate genomic events silencing FASTKD2 mRNA expression. The promoter of the FASKD2 gene contains a predicted CpG island suggesting a methylation-dependent silencing in *SLC30A10^Δ^* cells. We measured methylation of the FASTKD2 promoter and adjacent regions by EpiTyper, a MALDI-TOF mass spectrometry-based bisulfite sequencing method for quantitative and region-specific DNA methylation analysis (38, 39) (Fig. 2E). Approximately, 10-30% of each amplicon encompassing the CpG island and the 5’ region of the FASTKD2 gene were methylated in wild type cells as compared to 75-100% of the same loci in *SLC30A10^Δ^* cells (Fig. 2E). There was no methylation detected in the first FASTKD2 intron in both genotypes (Fig. 2E). The expression of MDH1B, a gene adjacent to FASTKD2 was low and remained unchanged in *SLC30A10^Δ^* cells (Supplementary table 3). These results demonstrate that the expression of FASTKD2 mRNA in *SLC30A10^Δ^* cells is prevented by methylation of its promoter and adjacent sequences.

### Components of the Mitochondrial RNA Granule are Downregulated after Manganese Homeostasis Disruption

FASTKD2 is part of the mitochondrial RNA granule required for mitochondrial RNA processing and ribosome assembly (26, 27, 40). FASTKD2 is engaged in protein complexes with other mitochondrial RNA binding proteins including DHX30, GRSF1, and mitochondrial ribosome subunits, such as MRPS18B (27, 41). Since FASTKD2 was the only common mitochondrial transcript downregulated in both *SLC30A10^Δ^* and *PARK2^Δ^* cells, we measured the abundance of FASTKD2, DHX30, GRSF1, and subunits of the mitochondrial ribosome by immunoblot. FASTKD2 protein was undetectable in *SLC30A10^Δ^* and *PARK2^Δ^* cells (Fig. 3A). Furthermore, the mitochondrial ribosome proteins MRPS18B and MRPL44 were both downregulated in both mutant cells, suggesting a perturbation of the mitochondrial RNA granule. To independently verify these results in neuronal cell types, we CRISPR edited the SLC30A10 gene in SH-SY5Y cells, a diploid neuroblastoma cell line (Fig. 3B). FASTKD2 and DHX30 expression were downregulated in a pool of cells where the SLC30A10 gene was acutely CRISPRed out in a 95% of all cells (Fig. 3B and S1). We next tested whether manganese exposure can induce this granule phenotype in iPCS-derived human astrocytes and glutamatergic neurons. Neurons were more sensitive to manganese with an IC50 of ~250 μM while astrocytes were resistant even after exposure to 1 mM manganese (Fig. 3C). Astrocytes were the most manganese resistant cells among all cell types analyzed (Fig. 3C and D). This increased astrocyte resistance correlated with SLC30A10 mRNA expression in single cell RNA seq experiments (Fig. 3E). Interestingly, iPSC-derived neurons downregulated the protein levels of DHX30, GRSF1, and MRPS18B when exposed to low manganese concentrations (Fig. 3F, 100 μM). Selection of *SLC30A10^Δ^* SH-SY5Y clones (Fig. S1B-C) recovered cells with decreased expression of FASTKD2, yet DHX30 and MRPS18B protein downregulation became evident after an acute manganese exposure of these mutant clones (Fig. S1E, compare lane 2 with 4 and 6).

**Figure 3.**
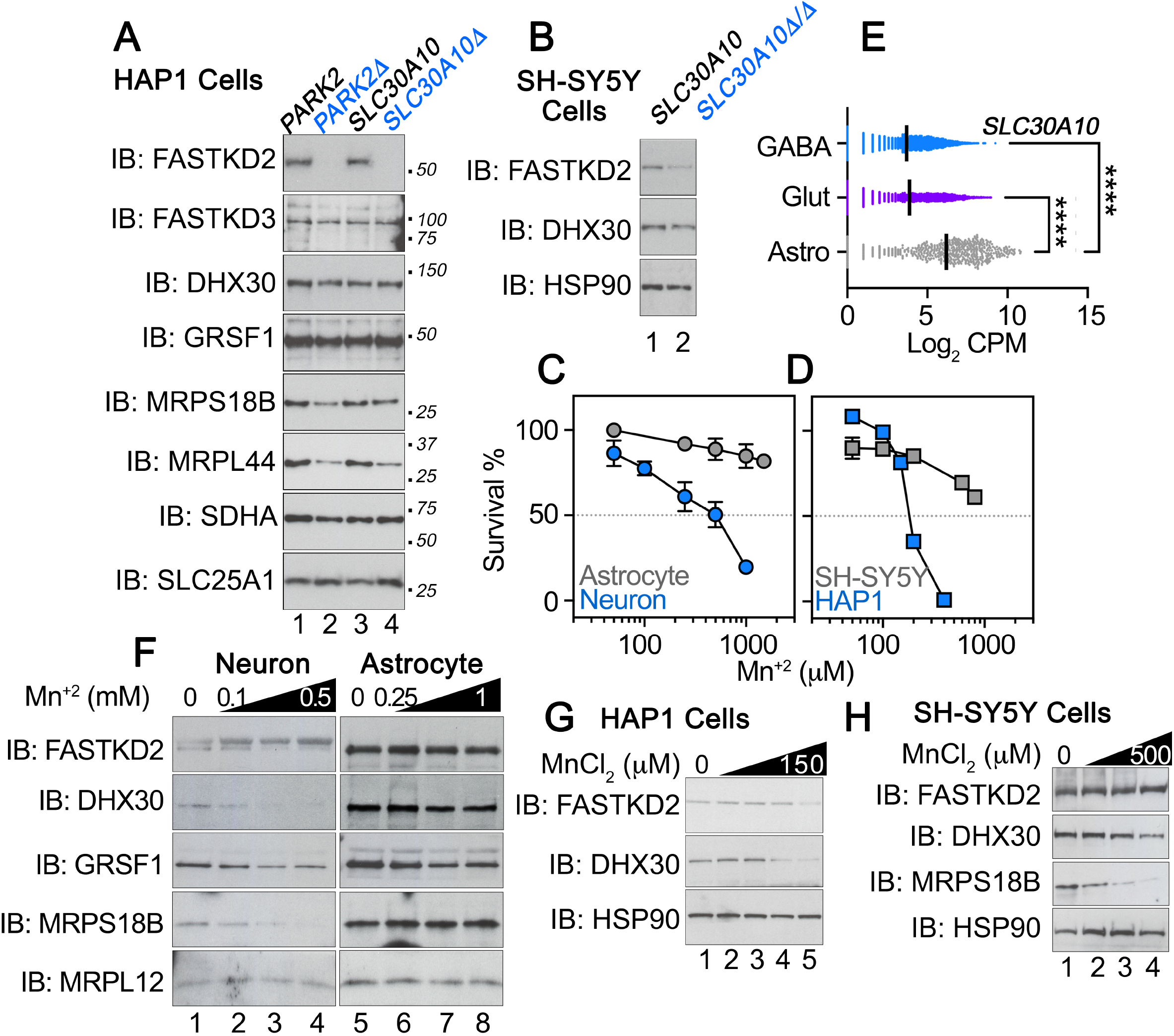
Downregulation of the Mitochondrial RNA Granule in SL30A10 Mutants and Cells Resistant to Manganese. **A.** Immunoblot analysis of wild type (odd lanes) and SLC30A10 or PARK2 mutant cells (even lanes). SDHA and SLC25A1, two mitochondrial proteins were used as loading controls. **B.** Cell extracts of wild type (lane 1) and a 95% CRISPR knock out SH-5YSY cell pool (lane 2) were analyzed by immunoblot with the indicated antibodies. HSP90 was used as a loading control. **C-D.** Comparative cell survival analysis of human iPSC-derived astrocytes and glutamatergic neurons differentiated for 15 days. HAP1 and SH-5YSY cells were simultaneously analyzed for comparison to iPSC-derived human cells. Cell survival in the absence of manganese correspond to 100%. Average ± SD is presented instead of SEM to make error bars visible, n=3. **E.** Expression of SLC30A10 mRNA in a single cell RNAseq dataset of mouse glutamatergic (n=8612, Glut) and GABAergic (n=3092, GABA) neurons as well as astrocytes (n=570, Astro). Kruskal-Wallis test followed by Benjamini-Hochberg q values <0.0001. **F.** iPSC-derived astrocytes and glutamatergic neurons (see C) incubated in the absence (lane 1) or presence of to increasing concentrations of manganese (lanes 2-4 and 6-8) were analyzed by immunoblot. The mitochondrial subunit MRPL12 was used as a loading control. **G**. HAP1 cells resistant to manganese were selected by exposure to increasing concentrations of manganese up to 150 μM for 5 days and analyzed at day 8 (lanes 2-5). Cell extracts were blotted with indicated antibodies. HSP90 was used as a loading control. **H.** SH-5YSY cells resistant to manganese were selected by exposure to increasing concentrations of manganese up to 500 μM for 5 days and analyzed at day 8 (lanes 2-4). Cell extracts were blotted with antibodies against indicated antigens. HSP90 was used as a loading control.

To distinguish whether these RNA granule phenotypes are associated with toxicity or are an adaptive mechanism, we exposed wild type HAP1 (Fig. 3G) and SH-SY5Y cells (Fig. 3H) to increasing concentrations of manganese for 9 doubling times to select manganese resistant cells. Manganese-resistant wild type HAP1 cells showed reduced levels of FASTKD2 and DHX30, indicating that manganese overload was sufficient to decrease expression of mitochondrial RNA granule components (Fig. 3G). Similarly, manganese resistant wild type SH-SY5Y cells downregulated DHX30 and MRPS18B (Fig. 3H). These results support a model where genetic and environmentally-induced manganese dyshomeostasis are sufficient to disrupt the composition and function of the mitochondrial RNA granule and that granule downregulation is an adaptive mechanism.

### Manganese Dyshomeostasis Disrupts the Function Of The Mitochondrial RNA Granule

The mitochondrial genome encodes 22 tRNAs, 13 mRNAs and 2 rRNAs expressed from two polycistronic transcripts (42–44). These polycistronic transcripts are processed by the mitochondrial RNA granule into mature forms of these three types of RNAs (43). Defects in mRNA precursor processing result in alterations in the expression of these RNAs and accumulation of unprocessed RNAs characterized by junctions between mRNAs-tRNAs, tRNAs-rRNAs, and between tRNAs (34, 43). Manganese-dependent downregulation of granule components predicts defective mitochondrial mRNA processing. Thus, we tested this model by analyzing the expression of mitochondria-encoded mRNAs by RNAseq and by NanoString with Mitostring, a panel of probes directed against unprocessed junctions (34).

We used the MitoString panel to quantify levels of unprocessed junctions in *SLC30A10^Δ^* and *PARK2^Δ^* mutant HAP1 cells and *SLC30A10^Δ/Δ^* SH-SY5Y cells. We found increased expression of tRNAs encoded in the mitochondrial genome in *SLC30A10^Δ^* and *PARK2^Δ^* mutant cells (Fig. 2A and 4A). *SLC30A10^Δ^* cells increased the expression of 14 tRNAs while *PARK2^Δ^* cells increased the expression of five tRNAs, all shared with *SLC30A10^Δ^* cells (Fig. 2A and 4A, TRNH, TRNL2, TRNM, TRNT, and TRNV). These changes in tRNA expression were independent from significant changes in the expression of most other mitochondrial RNA transcripts (Fig. 4A). We mapped the RNAseq reads for these tRNA to the mitochondrial genome and found increased expression of RNAs spanning tRNAs and their junctions to rRNAs yet without increased expression of the mitochondrial rRNAs RNR1 and RNR2 (Fig. 4B). The levels of the TRNF_RNR1 and TRND_COX2 junctions increased in *SLC30A10^Δ^* mutant cells, and these phenotypes were magnified when cells were exposed to manganese (Fig. 4C and S2). The same basal and manganese-induced outcomes were observed in two *SLC30A10 ^Δ/Δ^* SH-SY5Y cell clones when we probed for the TRNF_RNR1 junction (Fig. 4D and S2). However, genotype-dependent increases in the TRND_COX2 junction and other tRNA-mRNA junctions became evident after manganese treatment of wild type and *SLC30A10^Δ/Δ^* SH-SY5Y cell clones (Fig. 4D, F, and S2). *PARK2^Δ^* mutant HAP1 cells increased the TRND_COX2 junction after manganese exposure (Fig. 4E and S2) as well as other tRNA-mRNA junctions (Fig. 4F). Finally, we analyzed unprocessed junctions in HAP1 wild type cells selected for manganese resistance after chronic exposure to manganese (Fig. 4F-G, 2 weeks). Manganese resistant cells also increased the levels of TRNF_RNR1 and TRND_COX2 junctions as well as other tRNA-mRNA junctions (Fig. 4F-G and S2). These results demonstrate the acute and chronic manganese exposure or genetic defects increasing manganese sensitivity disrupt RNA processing by the mitochondrial RNA granule.

**Figure 4.**
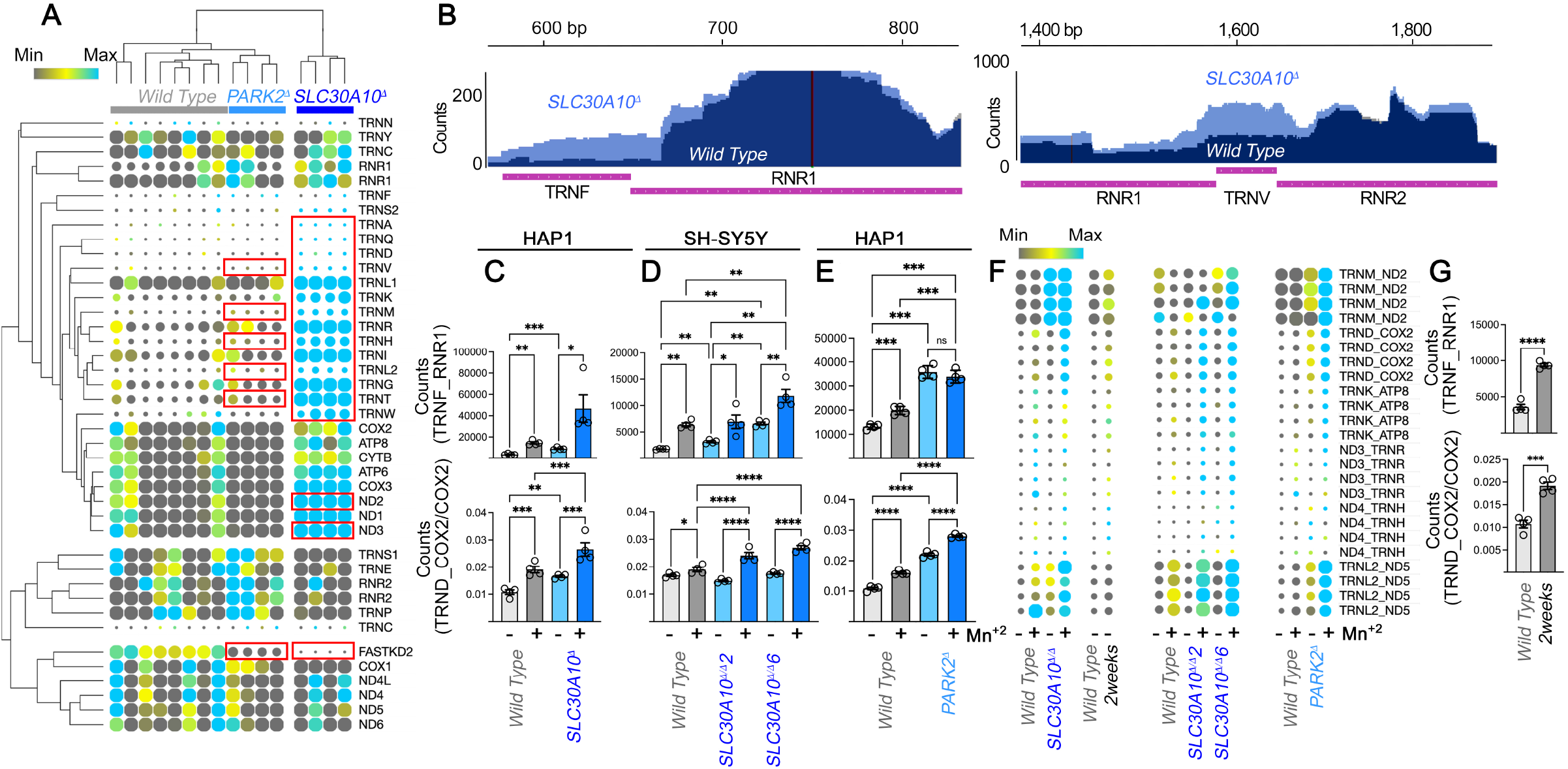
Processing of the Mitochondrial Transcriptome is Altered in SL30A10 and PARK2 Mutants. **A.** RNAseq analysis of mitochondrial encoded RNAs and the nuclear encoded FASTKD2 mRNA in wild type, SLC30A10 and PARK2 mutant HAP1 cells. Heat map represents (log_2_ expression CPM-row median/row median absolute deviation). Kendall Tau row and column clustering. Red boxes mark RNAs whose expression was significantly altered as determined by DESeq2 (p<0.05, Wald test corrected by Benjamini and Hochberg). **B.** Mapping of RNAseq reads to the reference human mitochondrial genome GRCh38. Depicted are examples of junctions between the tRNA TRNF and the RNR1 rRNA as well as the junctions flanking the tRNA TRNV. **C-G**. Nanostring analysis of the mitochondrial RNA transcriptome using the Mitostring panel. Probes detect unprocessed junctions and mitochondrial mRNAs. Data are expressed as a ratio of a junction to its corresponding mRNA except for the TRNF_RNR1 junction. Wild type cells and mutants were incubated overnight either in the presence or in the absence of a IC100 manganese dose, which achieved comparable intracellular levels of Mn (not shown). **C**. Data for wild type (300μM Mn and SLC30A10 mutant (150μM Mn) HAP1 cells. **D**. Data for wild type (1.6mM Mn) and two clones of SLC30A10 mutant (800μM Mn) SH-5YSYcells described in Figure S1. **E**. Data for wild type (300μM Mn) and PARK2 mutant (300μM Mn) HAP1 cells. **F**. Heat map represents of junction expression levels normalized to their corresponding mRNA (log_2_ expression CPM-row median/row median absolute deviation). Each data point is depicted and all these changes are significant. See below. **F**. Data for wild type cells selected for their resistance to manganese after two weeks of metal exposure. For C, D, E and G average ± SEM, n=4. One-Way ANOVA followed by Benjamini, Krieger and Yekutieli corrections. For G, unpaired t-test. Heat maps in A and F, symbol size is proportional to value.

### Electron Transport Chain Complexes are Disassembled in SLC30A10 Mutant Cells

Defects in mitochondrial RNA processing impair mitochondrial protein synthesis, which in turn results in downregulation of respiratory electron transport chain complex subunits (43). The *SLC30A10^Δ^* cell proteome uncovered downregulation of subunits in the four respiratory complexes whose assembly requires subunits synthesized by the mitochondrial ribosome (Fig. 5A, ***bold italic***). These findings predicted biochemical and functional defects in mitochondrial respiration and the assembly of electron transport chain complexes.

**Figure 5.**
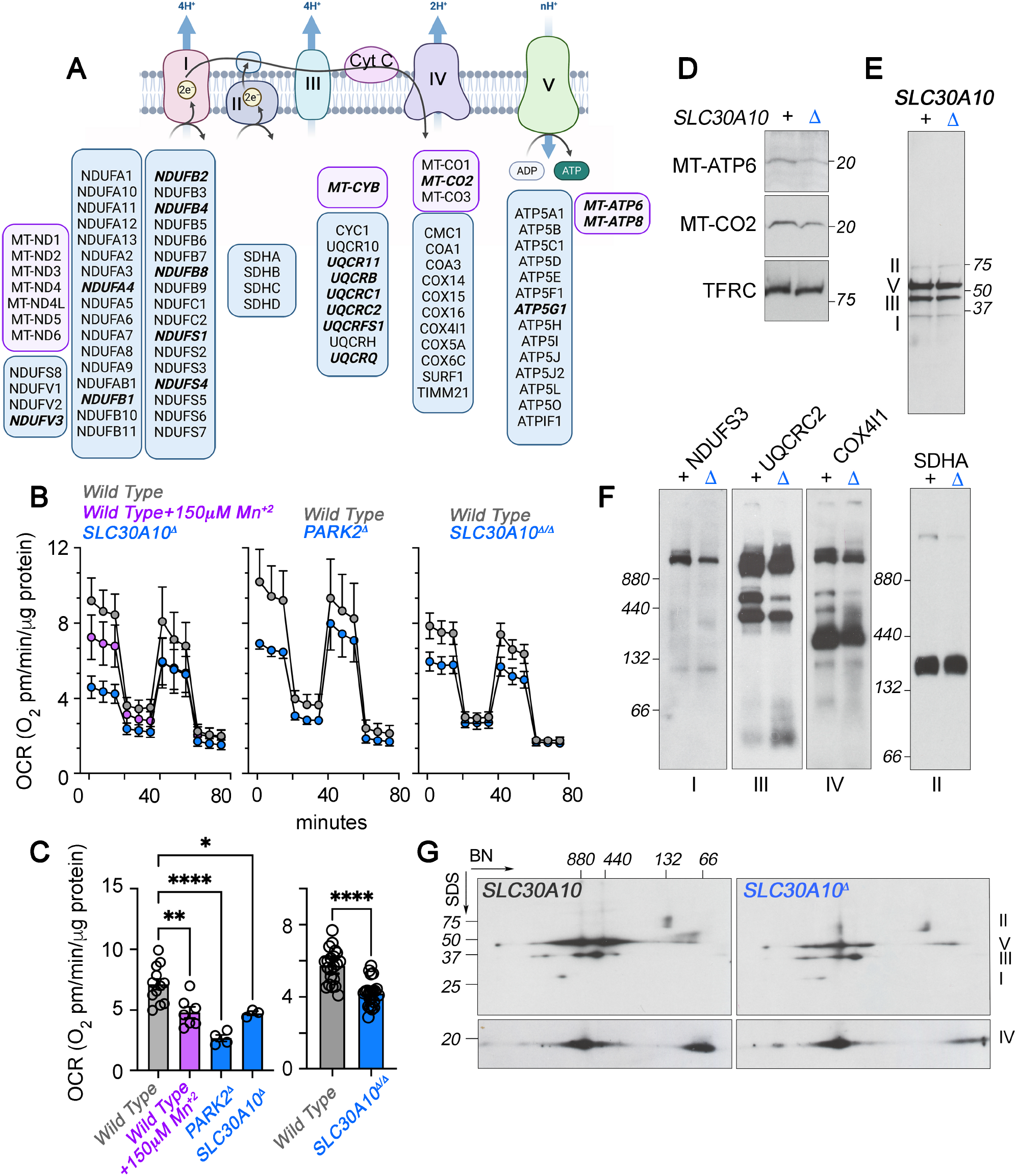
The Electron Transport Chain Assembly is Compromised in SLC30A10 Cells. **A.** Diagram of the electron transport chain complexes. Subunit enclosed in blue and purple rectangles are encoded in the nuclear and mitochondrial genomes, respectively. All bold italic subunits were downregulated in the proteome of SLC30A10 mutant cells. **B**. Stress test analysis of oxygen consumption rates in wild type, SLC30A10 and PARK2 mutant cells, and wild type cells treated overnight with manganese (left and middle panel show HAP1 cells). Right panel presents wild type and SLC30A10 mutant SH-5YSY cell pool. Average ± SEM. **C**. Basal respiration of cells described in B, average ± SEM. One-Way ANOVA followed by Dunnett corrections. **D**. Mitochondrial-encoded electron transport chain subunits are decreased in membrane fractions of SLC30A10 mutant cells. Transferrin receptor (TFRC) was used as a loading control in immunoblots. **E**. Expression levels of electron transport chain complexes I, III and V is reduced in mitochondrial fractions of SLC30A10 mutant cells. SDS-PAGE under reducing conditions. Immunoblotting antibodies are listed in F-G. **F-G**. Assembly of the electron transport chain is impaired in SLC30A10 mutant cells. **F**. Blue native electrophoresis (BN) of complexes I to IV in wild type and SLC30A10 mutant mitochondrial fractions. Complex II was used as a loading control in immunoblots. **G**. Two-dimensional electrophoresis of complexes I-V of the electron transport chain. Immunoblotting antibodies are listed in F except for the complex V antibody against ATP5A. All experiments in panels D-G were performed in HAP1 cells.

We measured mitochondrial respiration with Seahorse flow oximetry in cells where manganese dyshomeostasis was induced by metal exposure or by genetic defects in SLC30A10 and PARK2 (Fig. 5A-B) (45). Wild type HAP1 cells exposed to a manganese concentration equal to the IC50 (Fig. 1A and 3F) decreased mitochondrial respiration as determined by the basal respiration rate (Fig. 5A-B). A similar phenotype was observed in *SLC30A10^Δ^* and *PARK2^Δ^* mutant HAP1 cells, both mutations conferring increased sensitivity to manganese exposure (Fig. 5A and B). The reduced respiration phenotype was also found in a pool of SH-SY5Y cells where the SLC30A10 gene was CRISPRed out to an extent of 95% of all cells (Fig. 5A and B). We next measured the expression of respiratory complex proteins with antibodies against subunits encoded in the mitochondrial (Fig. 5D) and nuclear genome (Fig. 5E-G) in *SLC30A10^Δ^* HAP1 cells. Two mitochondrial-encoded proteins whose expression was down-regulated in the *SLC30A10^Δ^* HAP1 proteome, MT-CO2 (Fig. 5A, complex IV) and MT-ATP6 (Fig. 5A, complex V), were also found decreased by immunoblot (Fig. 5D). Concomitant with a reduction in these two mitochondrial-encoded proteins, we found decreased expression of the nuclear-encoded complex II subunit UQCRC2 (Fig. 5E, III) and the complex V subunit ATP5A (Fig. 5E, V), as well as the complex I subunit NDUFS3 (Fig. 5E, I). The expression of these subunits is sensitive to the assembly status of respiratory complexes. SDHA, a subunit of complex II, which is all encoded in the nuclear genome (Fig. 5A), was not modified in mass spectrometry and immunoblot experiments (Fig. 5A and E, II). Finally, we analyzed if decreased levels of these respiratory chain subunits affected the assembly of respiratory supercomplexes using blue native electrophoresis alone or in combination with SDS-PAGE (Fig. 5F-G). Complex I, III, IV, and V assembly into supercomplexes was compromised in *SLC30A10^Δ^* cells as determined by immunoblot of either blue native-, bidimensional-, or both electrophoreses (Fig. 5F-G). We did not detect modifications of complex II in both types of electrophoresis (Fig. 5F-G). We conclude that the function and assembly of the respiratory chain are impaired in *SLC30A10^Δ^* cells.

### Defects in mitochondrial RNA processing protect from a manganese challenge

The respiratory chain complex phenotypes observed in genetic and environmental manganese challenge models could be part of a pathogenesis mechanism or could represent cellular adaptive responses as suggested by the isolation of cells resistant to manganese (Fig. 3G-H). We discriminated between these two hypotheses using genetic and pharmacogenomic tools (Fig 6). We measured manganese sensitivity in wild type and mutant alleles of the FASTKD2 or DHX30 gene. We chose these genes as they are required for mitochondrial RNA granule function and because the expression of one or both of them was down-regulated in diverse forms of manganese dyshomeostasis (Fig. 3) (26–28, 34). *FASTKD2^Δ^* and *DHX30^Δ^* cells were significantly more resistant to manganese as compared to wild type cells (Fig 6A-B and Fig. S4C). We focused on the *FASTKD2^Δ^* cells to ascertain mechanisms accounting for this resistant phenotype (Fig. 6A). Two clones of *FASTKD2^Δ^* cells have complex changes in the expression of manganese efflux and influx transporters. For example, mRNA levels of two efflux transporters SLC30A10 and SLC40A1 mRNA were decreased to 23% and 63% in *FASTKD2^Δ^* cells as compared to a housekeeping gene VIM (Fig. 6C). However, we found that the expression of influx manganese transporters was each decreased to ~60% in these mutant cells (Fig. 6C, SLC11A2, SLC39A8 and SCL39A14) (46, 47). These transporter phenotypes were different from the phenotypes in *SLC30A10^Δ^* cells (Fig. 6C). This *FASTKD2^Δ^* transporter expression profile resulted in an attenuated capacity of these cells to store added manganese as measured by ICP mass spectrometry (Fig. 6E). However, neither transporter expression profiles nor the amount of accumulated metal, correlated with the relative resistance by each clone to manganese. These observations support our contention that loss of FASTKD2 in granule function is a contributing manganese resistance mechanism.

**Figure 6.**
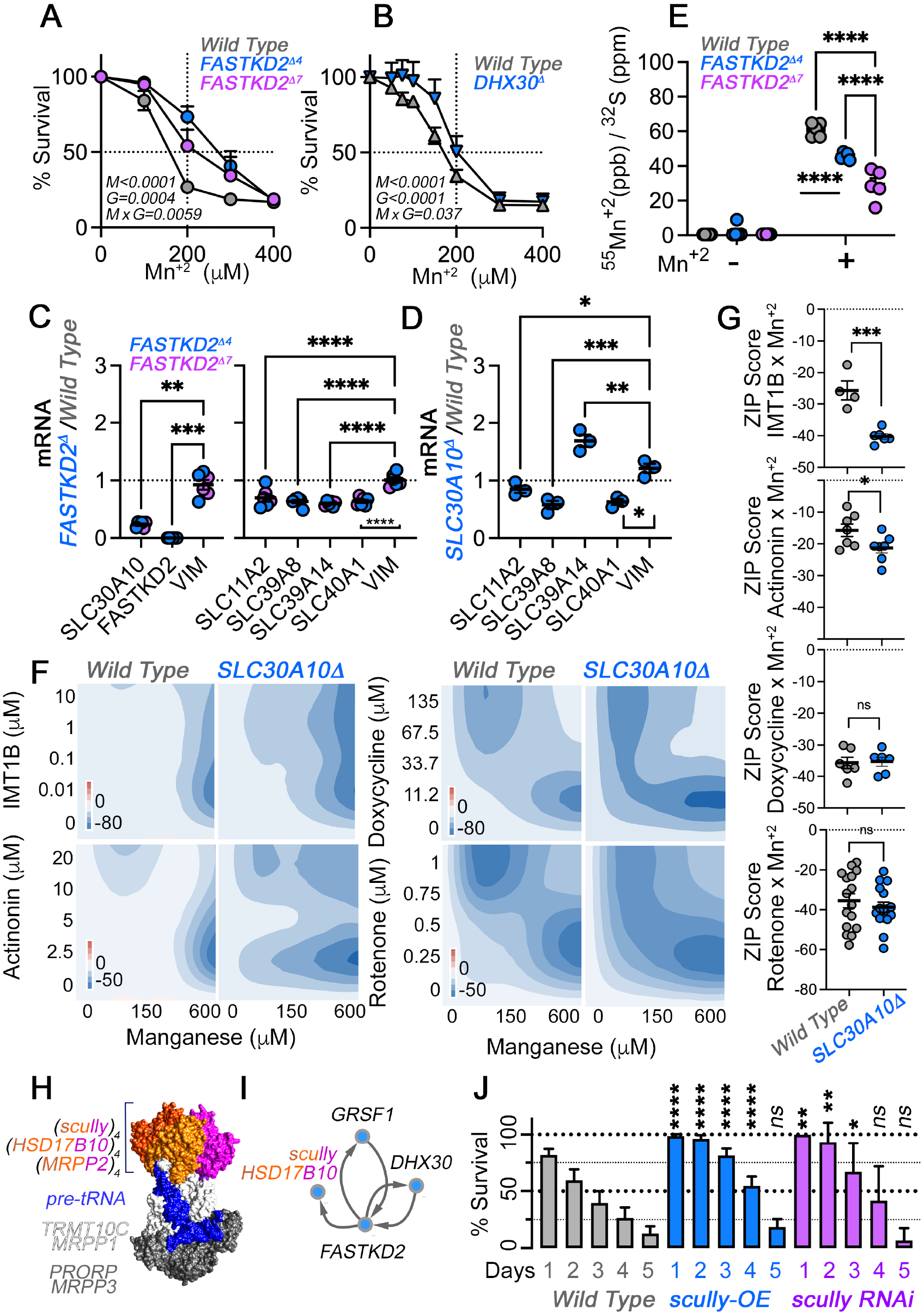
Defects in the Mitochondrial RNA Granule Protect Against Acute Manganese Toxicity. **A-B.** Cell survival analysis of with type, FASTKD2, and DHX30 mutant cells exposed to increasing concentrations of manganese. Two clones of FASTKD2 HAP1 mutant cells were tested. Average ± SEM, n=6 for FASTKD2 and their control cells and n=4 for the DHX30 mutant and its corresponding control. Two-Way ANOVA followed by Bonferroni corrections. **C-D**. mRNA levels of SLC30A10 and other manganese transporters in FASTKD2 and SLC30A10 mutant HAP1 cells. Vimentin mRNA (VIM) was used as loading control. Average ± SEM, for C n=6 (each clone is color coded, see legend) and for D n=3, Two-Way ANOVA followed by Šydák’s multiple corrections. **E**. ICP mass spectrometry determination of manganese in wild type and FASTKD2 mutant HAP1 cells treated with vehicle or manganese overnight. Data were normalized to total sulfur content. Average ± SEM, n=5, Two-Way ANOVA followed by Bonferroni corrections. **F**. Manganese-Drug interaction Synergy map using the Zero Interaction Potency (ZIP) score for cell survival. 26949479. Scores <-10 indicate an antagonistic interaction between metal and drug. Maps were generated with at least 6 independent experiments per metal-drug pair. **G**. Average ZIP score for manganese-drug interactions in wild type and SLC30A10 mutant cells. Average ± SEM, unpaired t-test. **H**. Structure of the human RNAse P complex. **I**. Proximity interactions between the RNAse P subunit *scully* (HSD17B10) and other mitochondrial RNA granule components from the mitochondrial high density proximity interaction map (41). **J**. Survival of adult Drosophila exposed to dietary manganese for 5 consecutive days in wild type, *scully* RNAi, or *scully* overexpressing animals (a dominant negative). Average ± 95% CI, see Fig. S5 for number of experiments and number of animals analyzed. One-Way ANOVA followed by Bonferroni corrections.

We further tested the protective effect of impaired RNA granule function by quantitative pharmacogenomic interaction experiments. We measured interactions between the SLC30A10 genotype, manganese, and drugs that disrupt mitochondrial protein synthesis from transcription to translation. We reasoned that much like *FASTKD2^Δ^* and *DHX30^Δ^* cells, pharmacological disruption of mitochondrial protein synthesis could be protective for manganese exposed cells. We used the Zero Interaction Potency model to quantitative assess synergistic or antagonistic effects of drugs on manganese-induced cell mortality (48, 49). We used IMT1B, an inhibitor of the mitochondrial RNA polymerase (50); actinonin, an agent that induces degradation of mitochondrial rRNAs and mRNAs (51); and doxycycline, a mitochondrial ribosome protein synthesis inhibitor (Fig. 6F-G) (52). All drugs antagonized the mortality induced by increasing concentrations of manganese both in both wild type and *SLC30A10^Δ^* cells as determined by Zero Interaction Potency scores under −10 (Fig. 6F-G, ZIP). Moreover, the antagonistic effects of IMT1B and actinonin were more pronounced in in *SLC30A10^Δ^* cells as compared to controls (Fig. 6F), a phenotype indicated by significantly lower ZIP scores in mutant cells (Fig. 6G). Doxycycline was the most effective antagonist of manganese-induced mortality and its effect was not modified by genotype (Fig. 6G), likely because of a ceiling effect. We conclude that disruption of the mitochondrial protein synthesis from transcription to translation increases the fitness of cells exposed to manganese.

We determined the organismal impact of disrupting mitochondrial RNA processing on manganese lethality by exposing adult *Drosophila* to dietary manganese. We genetically modified the function of the mitochondrial RNAse P, a complex required for the processing of the 5’ end of mitochondrial tRNAs, which is present in the RNA granule (Fig. 6H-I) (27, 41, 53, 54). One of the RNAse P subunits is encoded by the gene *scully* in *Drosophila. Scully* or its human orthologue, HSD17B10, is present as a tetramer in the RNAse P complex (Fig. 6H). Furthermore, HSD17B10 is associates with FASTKD2 and DHX30 in human mitochondria (Fig. 6I). *Scully* overexpression or RNAi impair mitochondrial function and tRNA processing (55, 56), thus, we tested the effects of *scully* up- or downregulation with the UAS-Gal4 system on animal survival after manganese exposure. UAS transgenes were expressed using a ubiquitous actin-GAL4 driver. Mortality was 20% after one day of manganese exposure and increased to 87% after five days of exposure in wild type animals (Fig. 6J). In contrast, *scully* overexpression, which acts as a dominant negative, or *scully* RNAi abolished manganese lethality after one day exposure (Fig. 6J). The protective effects of *scully* manipulations remained for three days (Fig. 6J). However, *scully*-mediated protection was indistinguishable from wild type animals at five days of dietary exposure. These results show that defects in mitochondrial RNA processing are adaptive for early stages of toxic manganese exposure at the organism level.

## Discussion

We identified cellular and molecular mechanisms altered by mutations in SLC30A10 and PARK2, both conferring manganese susceptibility. We then used these mechanisms to ask how disruption of these processes would impact cell and animal fitness after a toxic manganese challenge. We investigated a mechanistic overlap between these genetic forms of manganism and Parkinson’s disease and found that these two genotypes result in a defective mitochondrial RNA granule. This granule impairment is driven by decreased expression of FASTKD2 and its interactors DHX30 and MRPS18B in both SLC30A10- and PARK2-null cells. Furthermore, we demonstrate a mitochondrial RNA granule function defect by alterations in the mitochondrial transcriptome and RNA processing using NanoString technology in SLC30A10 and PARK2 mutant cells as well as cells exposed to manganese irrespective of their genotype. The mitochondrial RNA granule is required for polycistronic mitochondrial transcripts processing into mature transfer, messenger and ribosomal RNA; thus, it is necessary for mitochondrial protein synthesis and the assembly of the electron transport chain (43, 54). Importantly, much like manganese-induced responses (57, 58), we find that SLC30A10- and PARK2-null cells have altered expression of nuclear and mitochondrial-encoded electron transport chain proteins, thus resulting in impaired respiratory function chain.

Even though it has been suggested that loss of electron transport chain function is non-adaptive/pathogenic in manganese exposure, this proposition has not been directly tested (reviewed in (59–61)). Furthermore, a non-adaptive/pathogenic model does not distinguish between short and long-term consequences of a defective electron transport chain after manganese exposure. We directly tested whether fitness to a manganese challenge was enhanced or decreased after loss-of-function mutations in mitochondrial RNA granule components or after selective inhibition of mitochondrial transcription-translation. All these experimental conditions ultimately impair the electron transport chain. To our surprise, our data demonstrate that decreased function of the mitochondrial RNA granule increases fitness after acute manganese exposure both in cells and adult *Drosophila*. This adaptive mechanistic proposition is robustly supported by multipronged approaches. First, editing of either FASTKD2 or DHX30 genes by CRISPR antagonized manganese-induced cell death using two cell survival assays. Second, wild type cells selected by their resistance to prolonged manganese exposure show downregulation of mitochondrial RNA granule protein expression and function. Third, inhibition of mitochondrial transcription, which resides upstream of the mitochondrial RNA granule, or inhibiting processes downstream the granule, such as mitochondrial protein synthesis or complex I function, results in increased survival to a manganese challenge, both in wild type and mutant cell. Collectively, this evidence demonstrates that impaired mitochondrial RNA granule function is an adaptive mechanism increasing resistance to manganese in a cell autonomous manner. Finally, we demonstrate that impairing the processing of mitochondrial tRNAs in adult *Drosophila*, either by overexpression or RNAi of the RNAse P subunit *scully*, acutely protects animals from manganese exposure. These findings are incompatible with a model where impairments in mitochondrial RNA granule function and downstream mechanisms decrease cellular and animal fitness during acute metal exposure. We propose that decrease mitochondrial respiration is an acute response to evade immediate cell and organismal death induced by toxicants. While this proposition may seem at odds with current models (59, 60), we think we have captured an early and upstream step of the existing model. We see our findings as expanding current models in manganism. The current conception is that manganese neurotoxicity in animal models is mediated by apoptosis due to manganese accumulation in mitochondria (60, 62, 63). Mitochondrial manganese would then result in disruption of respiratory chain function with induction of oxidative stress, transition pore opening and caspases release (60, 64). In this model, mechanisms causative of reactive oxygen species production and apoptosis are unclear as they have been attributed to multiple putative manganese targets including complex I and II, and mitochondrial matrix oxidases (64–66). We think that the disruption of the mitochondrial RNA granule can account for many of these manganese-dependent mitochondrial phenotypes affecting the respiratory chain and redox metabolism in cells. We argue that such defects are protective in acute manganese exposure conditions yet deleterious and protracted during chronic manganese exposure. Our Drosophila experiments support this idea. Impairing tRNA processing in whole animals is protective in the first three days of manganese dietary exposure. However, a prolonged exposure to manganese for 5 days is equally lethal to wild type and mutant flies. We propose that chronic mitochondrial dysfunction would pave the way for neuronal cell death. Our model borrows from recent evidence with a conditional allele of complex I in dopaminergic cells that causes Parkinson’s disease (67). In this Parkinson’s mouse model, electron chain disruption causes a rewiring of neuronal metabolism to aerobic glycolysis without cell death in early stages. Only after prolonged mitochondrial dysfunction of the electron transport chain neuronal cell death becomes a dominant trait (67). This deleterious outcome of chronic perturbation of oxidative phosphorylation may be a universal consequence of rewiring the bioenergetic metabolism away from the electron transport chain (68).

Our results demonstrate RNA granule dysregulation as a mechanism affected by manganese dyshomeostasis. However, our findings also suggest cell type- and genotype-dependent variations in the way mitochondrial granule components respond to manganese exposure. We uncovered one mechanism dominated by complete downregulation of FASTKD2 expression in SLC30A10-null HAP1 cells. In these mutant cells, FASTKD2 expression was abrogated by methylation of the FASTKD2 promoter, an event likely facilitated by the near-haploid nature of the genome in HAP1 cells (69). However, we found that FASTKD2 expression was only partially decreased or remained unaltered, in SH-SY5Y cells and iPSC-derived human neurons, respectively. Instead in these cells, granule components and FASTKD2 interactors such as DHX30, GRSF1, and MRPS18B were affected either by SLC30A10 deficiency or by manganese exposure. Furthermore, we found that manganese exposure changed the SDS-PAGE migration of FASTKD2 in iPSC-derived human neurons but not in other cell types, suggesting cell type specific post-translational modifications induced by manganese. These cell-type variations were also evident in the quality of the unprocessed mitochondrial RNA intermediaries accumulated across cell type, exposure status, and genotype. We speculate that these findings reflect diversified and complex pathways impinging on mitochondrial RNA processing in different cell types. These arguments about the complexity of mitochondrial RNA processing mechanisms are reinforced by loss-of-function screening of mitochondrial RNA binding proteins (34). This screen has revealed that loss-of-function of SSBP1 increases the content of RNAs encoded on the light strand of the mitochondrial genome (34). However, we found that despite increased SSBP1 levels in SLC30A10 mutant cells, we still observed increased content of transcripts encoded by the light strand Finally, our findings demonstrate that PARK2-dependent pathways also control the composition of the granule and its function. However, the profile of mitochondrial RNA processing intermediaries whose content increased only partially overlaps with SLC30A10 deficient cells even though these mutations were engineered in an isogenic cell system. We postulate that multiple trajectories converge on the mitochondrial RNA processing machinery to modulate its function in normal and toxicant-exposed conditions.

## Materials and Methods

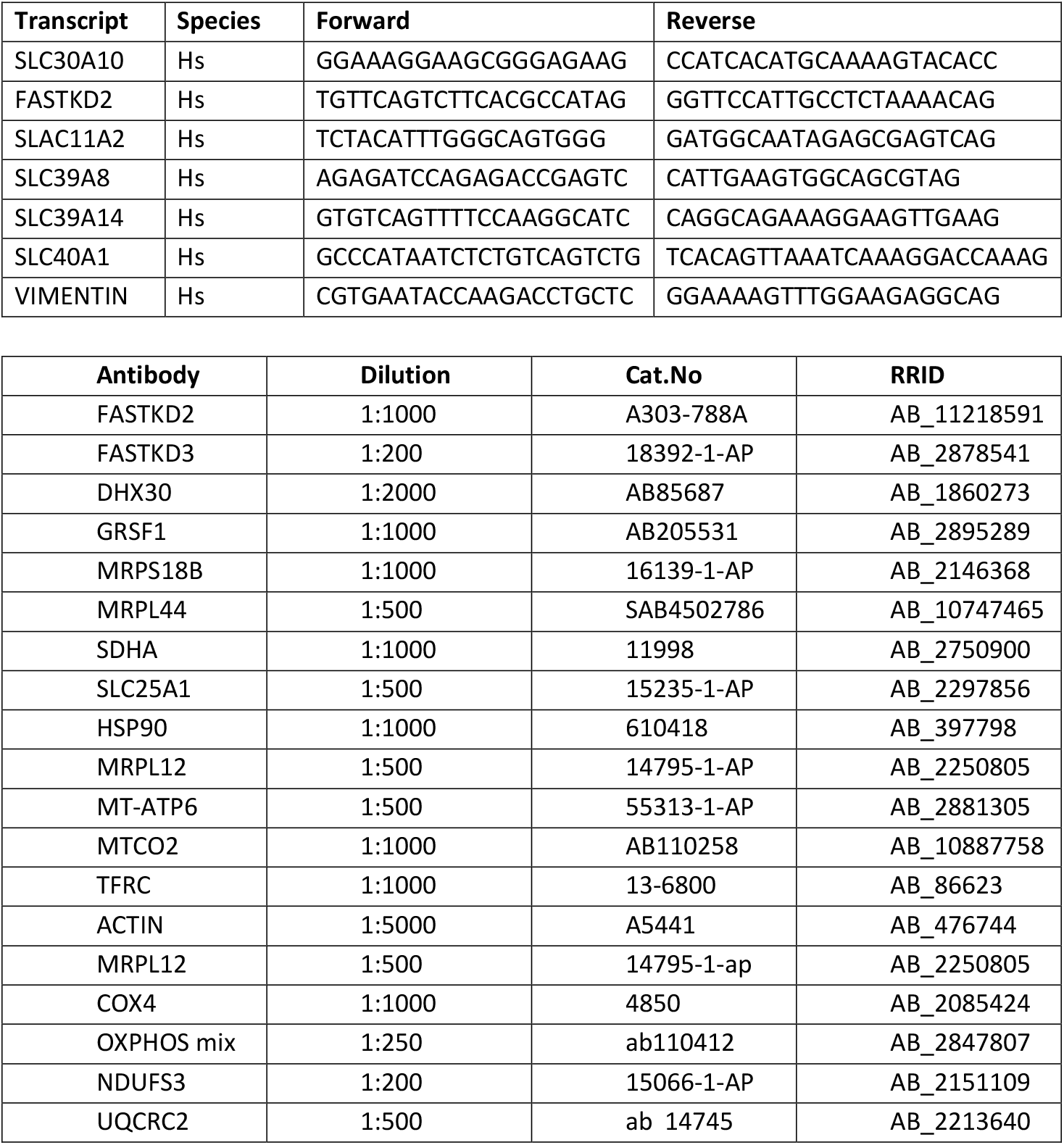

### Cell Lines and Culture Conditions

Human haploid (HAP1) isogenic cell lines were obtained from Horizon Discovery. The cells were either wild type HAP1 (RRID: CVCL_Y019) or gene-edited for SLC30A10 (HZGHC004693c005, RRID:CVCL_TM87), PARK2 (HZGHC003208c002, RRID:CVCL_TC07), SLC25A4 (HZGHC000778c011, RRID:CVCL_TM45), FASTKD2 (HZGHC006859c004, HZGHC006859c007 RRID CVCL_B5J3 AND CVCL_B5J4) or DHX30 (HZGHC007945c004 RRID: CVCL_B5J6). Cell lines were maintained in IMDM media (Lonza Walkersville, 12-722F) supplemented with 10% FBS and 100 μg/ml penicillin and streptomycin. The cell cultures were grown at 37°C in a 10% CO_2_ incubator. Manganese resistant HAP1 cells were obtained by incubating wild type Hap cells in 150 μM Manganese until cell death ceased, for 5 days and then grown in regular media. Human neuroblastoma SH-SY5Y cells (ATCC, CRL-2266; RRID:CVCL_0019) were grown in DMEM media containing 10% FBS at 37°C in 10% CO_2_. SH-SY5Y deficient in SLC30A10 were genome edited using gRNA and Cas9 preassembled complexes by Synthego with a knock-out efficiency of 98%. The gRNAs used was GCAGCGCGAUGGAGUUGCCC which targeted transcript ENST00000366926.3 exon 1. Wild-type and mutant cells were cloned by clonal dilution and mutagenesis was confirmed by Sanger sequencing with the primer: 5’GAGACAATCTGGGAGGCGG.

### Generation of iNeurons from Human iPSCs

iPSCs were dissociated with accutase (Gibco, A11105) and plated at 380,000 cells per well of a 12-well plate on day −2. Cells were plated on matrigel (Corning, 354230)-coated plates in mTeSR medium (StemCell Technologies, 85850). On day −1, *hNGN2* lentivirus (TetO-h*NGN2*-P2A-PuroR (Addgene, 79049) or TetO-h*NGN2*-P2A-eGFP-T2A-PuroR (Addgene, 79823)) together with FUdeltaGW-rtTA (Addgene, 19780) lentivirus at 1×10^6^ pfu/ml per plasmid (multiplicity of infection (MOI) of 2) were added in fresh mTeSR medium containing 4μg/μl polybrene (Sigma, TR-1003). On the day 0, the culture medium was replaced with fresh KSR containing KnockOut DMEM (Gibco, 10829018), Knockout Replacement Serum (Gibco, 10828028), 1X Glutamax (Gibco, 35050061), 1 X MEM Non-essential Amino Acids (NEAA, Gibco, 11140050) and 100μM β-Mercaptoethanol (Gibco, 21985023). Doxycycline (2μg/ml) was added on day 0 to induce TetO gene expression and retained in the medium until the end of the experiment. On day 1, puromycin selection (5μg/ml) period was started. On day 4, the culture medium was replaced by neuron medium: Neurobasal medium (Gibco, 21103049) supplemented with B27 supplement (Gibco, 17504044), 1 X Glutamax, 20% Dextrose (Sigma,D9434) containing 10ng/ml BDNF (Peprotech, 45002), 10ng/ml GDNF (PeproTech, 45010), 2μg/ml doxycycline (Sigma, D9891) and 5μg/mL puromycin (InvivoGen, ant-pr-1). Beginning at day 7, half of the medium in each well was replaced every week. h*NGN2*-induced neurons were assayed on day 24 (70).

### Differentiation of iPSCs into forebrain specific neural progenitors and astrocyte

iPSCs cell colonies were detached with 1 mg/ml collagenase (Therm Fisher Scientific, 17104019) treatment for 1 h and suspended in embryoid body (EB) medium, consisting of DMEDM/F12 (Gibco, 11330032), 20% Knockout Serum Replacement (Gibo, 10828028), 1 X Glutamax (Gibco, 35050061), 1 X MEM Non-essential Amino Acids (NEAA, Gibco, 11140050), 100μM β-Mercaptoethanol (Gibco, 21985023), 2 μM dorsomorphin (Tocris, 3093) and 2 μM A-83 (Tocris, 692), in non-treated polystyrene plates for 7 days with a daily medium change. After 7 days, EB medium was replaced by neural induction medium (hNPC medium) consisting of DMEM/F12, 1 X N2 supplement (Gibco, 17502048), B27 supplement, 1X Non-essential Amino Acids, 1 X Glutamax, 2 μg/ml heparin (Sigma) and 2 μM cyclopamine (Tocris, 1623). The floating EBs were then transferred to Matrigel-coated 6-well plates at day 7 to form neural tube-like rosettes. The attached rosettes were kept for 15 days with hNPC medium change every other day. On day 22, the rosettes were picked mechanically and transferred to low attachment plates (Corning) in hNPC medium containing B27.

For astrocytes differentiation (71), resuspended neural progenitor spheres were dissociated with accutase at 37°C for 10 min and placed on Matrigel coated 6 well plate. Forebrain NPCs were maintained at high density in hNPC medium. Forebrain NPCs were differentiated to astrocytes by seeding dissociated single cells at 15,000 cells/cm^2^ density on Matrigel-coated plates in astrocyte medium (ScienCell: 1801, astrocyte medium (1801-b), 2% fetal bovine serum (0010), astrocyte growth supplement (1852) and 10U/ml penicillin/streptomycin solution (0503)). From day 2, cells were fed every 48 hours for 20-30 days. When the cells reached 90-95% confluency (approximately every 6-7 days), they were split to the initial seeding density (15,000 cells/cm^2^) as single cells in astrocyte medium

### Immunoblotting

Cells were grown in 10 cm or 15 cm dishes up to required confluency. The plates were placed on ice, and the cells were washed with cold PBS (Corning, 21-040-CV) containing 0.1 mM CaCl_2_ and 1.0 mM MgCl_2_. Lysis buffer containing 150 mM NaCl, 10 mM HEPES, 1 mM EGTA, and 0.1 mM MgCl_2_, pH 7.4 (Buffer A) with 0.5% Triton X-100 and Complete anti-protease (Roche, 11245200) was added to each plate. Cells were then scraped and placed in Eppendorf tubes on ice for 30 min and centrifuged at 16,100 × *g* for 10 min. The insoluble pellet was discarded and the clarified supernatant was recovered. Bradford Assay (Bio-Rad, 5000006) was used to determine protein concentration and all lysates were flash frozen on dry ice and stored at −80°C.

Cell lysates were reduced and denatured with Laemmli buffer (SDS and 2-mercaptoethanol) and heated for 5 min at 75°C. Equivalent amounts of samples were loaded onto 4%-20% Criterion gels (Bio-Rad, 5671094) for SDS-PAGE and transferred using the semidry transfer method to PVDF membranes (Millipore, IPFL00010). The PVDF membranes were incubated in TBS containing 5% nonfat milk and 0.05% Triton X-100 (TBST; blocking solution) for 30 min at room temperature. The membrane was then rinsed thoroughly and incubated overnight with optimally diluted primary antibody in a buffer containing PBS with 3% BSA and 0.2% sodium azide. The next day, membranes were rinsed in TBST and treated with HRP-conjugated secondary antibodies against mouse or rabbit (ThermoFisher Scientific, A10668; RRID:AB_2534058; RRID:AB_2536530) diluted 1:5000 in the blocking solution for at least 30 min at room temperature. The membranes were washed in TBST at least 3 times and probed with Western Lightning Plus ECL reagent (PerkinElmer, NEL105001EA) and exposed to GE Healthcare Hyperfilm ECL (28906839).

### Total RNA extraction, cDNA preparation, and qRT-PCR

Cells were grown on 10 cm plates and total RNA was extracted using the Trizol reagent (Invitrogen, 15596026). Cells were rinsed twice in PBS containing 0.1 mM CaCl_2_ and 1.0 mM MgCl_2_, followed by 1 ml of Trizol added to each plate. Cells were scraped and collected in an Eppendorf tube and incubated for 10 min at room temperature on an end-to-end rotator. 200 μl of chloroform was then added to each tube; and after a 3 minute incubation, the mixture was centrifuged at 12,000 RPM for 15 min at 4°C. The colorless aqueous layer was carefully collected and 500 μl of isopropanol was added to it. This mixture was rotated for 10 min at room temperature followed by centrifugation at 12,000 rpm for 15 min. The pellet was washed with 75% ethanol after discarding the supernatant. Air-dried pellets were dissolved in 20-100 μl of molecular grade RNAase-free water. RNA was quantified and the purity determined using the Nanodrop One^C^ (Thermo Fisher Scientific) and the samples were diluted to 1 μg/μl.

Superscript III First Strand Synthesis System Kit (Invitrogen, 18080-051) and random hexamer primers were used to synthesize cDNA. 5 μg RNA was incubated with the hexamers, dNTPs at 65°C for 5 min and then placed on ice. cDNA synthesis mix was added to each of the tubes and the samples underwent the following protocol: 25°C for 10 min followed by 50 min at 50°C. The reaction was terminated 85°C for 5 min and the samples treated with RNase H at 37°C for 20 mins. The synthesized cDNA was stored at −20°C. IDT Real-Time qPCR Assay Entry site (https://www.idtdna.com/scitools/Applications/RealTimePCR/) was used to design primers. Primers were obtained from Sigma-Aldrich Custom DNA Oligo service. To control for primer quality and specificity, we used primer annealing and melting curve parameters. The primer list is provided in table above. qRT-PCR was performed with 1μl of the synthesized cDNA using LightCycler 480 SYBR Green I Master (Roche, 04707516001) on a QuantStudio 6 Flex instrument (Applied Biosystems) in a 96 well format. Initial denaturation was performed at 95°C for 5 min, followed by 45 cycles of amplification with a 5 s hold at 95°C ramped at 4.4°C/s to 55°C. Temperature is maintained for 10 s at 55°C and ramped up to 72°C at 2.2°C/s. Temperature was held at 72°C for 20 s where a single acquisition point was collected and then ramped at 4.4°C/s to begin the cycle anew. The temperature was then held at 65° for 1 min and ramped to 97°C at a rate of 0.11°C/s. Five acquisition points were collected per °C. Standard curves were used for quantification for each primer set using the QuantStudio RT-PCR Software version 1.2. Data were standardized using housekeeping genes and presented as a ratio of control to experimental samples.

### TMT Mass Spectrometry

Samples were washed twice in 1X PBS and lysed in 8M urea, 50mM Tris HCl, pH 8.0, 1X Roche Complete Protease Inhibitor and 1X Roche PhosStop in a Next Advance Bullet Blender using 1.0mm silica beads in. Lysates were quantified by Qubit fluorometry (Life Technologies). 30μg of each sample was digested overnight with trypsin. Briefly, samples were reduced for 1h at RT in 12mM DTT followed by alkylation for 1h at RT in 15 mM iodoacetamide. Trypsin was added to an enzyme:substrate ratio of 1:20. Each sample was acidified in formic acid and subjected to SPE on an Empore SD C18 plate (3M catalogue# 6015 SD). Each sample was lyophilized and reconstituted in 140mM HEPES, pH 8.0, 30% acetonitrile for TMT labeling. 40μL of acetonitrile was added to each TMT tag tube and mixed aggressively. Tags were incubated at room temperature for 15min. 15μL of label was added to each peptide sample and mixed aggressively. Samples were incubated in an Eppendorf Thermomixer at 300rpm 25°C for 1.5h. Reactions were terminated with the addition of 8μL of fresh 5% hydroxylamine solution and 15 min incubation at room temperature. Each labeled sample was pooled into two experiments, frozen, and lyophilized and subjected to SPE on a High-Density 3M Empore SDB-XC column (Cat. #4340-HD). The eluent was lyophilized. The 300μg per 10-plex was subjected to high pH reverse phase fractionation as follows:Buffer A: 10mM NaOH, pH 10.5, in water Buffer B: 10mM NaOH, pH 10.5, acetonitrile. Samples were run on a XBridge C18 column, 2.1mm ID x 150mm length, 3.5μm particle size (Waters, part #186003023). We used a HPLC system: Agilent 1100 equipped with a 150μL sample loop operating at 0.3mL/min, detector set at 214nm wavelength. High pH Reverse Phase Fractionation: 300μg of dried peptide was resolubilized in 150μL of Buffer A and injected manually. Fractions were collected every 30s from 1-49min (96 fractions total, 150μL/fraction). Peptides (10% per pool) were analyzed by nano LC/MS/MS with a Waters NanoAcquity HPLC system interfaced to a ThermoFisher Fusion Lumos mass spectrometer. Peptides were loaded on a trapping column and eluted over a 75μm analytical column at 350nL/min; both columns were packed with Luna C18 resin (Phenomenex). Each high pH RP fraction was separated over a 2h gradient (24h instrument time total). The mass spectrometer was operated in data-dependent mode, with MS and MS/MS performed in the Orbitrap at 60,000 FWHM resolution and 50,000 FWHM resolution, respectively. A 3s cycle time was employed for all steps. Data Processing Data were processed through the MaxQuant software v1.6.0.16 (www.maxquant.org) which served several functions: 1. Recalibration of MS data. 2. Filtering of database search results at the 1% protein and peptide false discovery rate (FDR). 3. Calculation of reporter ion intensities (TMT). 4. Isotopic correction of reporter ion intensities (TMT). Data were searched using Andromeda with the following parameters: Enzyme: Trypsin Database: Swissprot Human Fixed modification: Carbamidomethyl (C) Variable modifications: Oxidation (M), Acetyl (Protein N-term) Fragment Mass Tolerance: 20ppm Pertinent MaxQuant settings were: Peptide FDR 0.01 Protein FDR 0.01 Min. peptide Length 7 Min. razor + unique peptides 1 Min. unique peptides 0 Second Peptides FALSE Match Between Runs FALSE The proteinGroups.txt file was uploaded to Perseus v1.5.5.3 for data processing and analysis.

### ICP MS

Equal number of cells were plated on 10 cm dishes and placed in the incubator at 37°C and 10% CO_2_. After a minimum of 16 hours the media on the plates was replaced with media containing requisite concentration of manganese chloride and as a control, plain media. Cells were then placed back in the incubator and grown overnight. On the day of sample collection, the plates were placed on ice and washed three times with PBS containing 10 mM EDTA (Sigma-Aldrich, 150-38-9). After the final wash, the cells were incubated with10ml of PBS and 10 mM EDTA for 20 min on ice. Cells were collected with mechanical agitation with a pipette and placed in a 15 ml falcon tube. Cells were pelleted at 800 × g for 5 min at 4°C. After discarding the supernatant, the cell pellet was washed with ice-cold PBS and the resuspended cells were centrifuged at 16,100 ×g for 5 min. The supernatant was aspirated and the residual pellet was immediately frozen on dry ice for at least 5 min and stored at −80°C for future use.

Cell pellets were digested by adding 20μL of concentrated (70%) trace metal basis grade nitric acid (Millipore Sigma) followed by heating at 95 °C for 10 minutes. After cooling samples were diluted to 800μL with 2% nitric acid (v/v). Metal levels were quantified using a triple quad ICP-MS instrument (ThermoFischer, iCAP-TQ) operating in oxygen mode under standard conditions (RF power 1550 W, sample depth 5.0 mm, nebulizer flow 1.12 L*min^-1^, spray chamber 3°C, extraction lens 1,2 −195, −215V). Oxygen was used as a reaction gas (0.3 ml*min^-1^) to remove polyatomic interferences or mass shift target elements (Analytes measured; 32S.16O, 55Mn, 45Sc.16O). External calibration curves were generated using multi-elemental standard (ICP-MS-CAL2-1, AccuStandard, USA) and ranged from 0.5-1000 μg*L^-1^ for each element. Scandium (10 μg*L^-1^) was used as internal standards and diluted into the sample in-line. Samples were introduced into the ICP-MS using the 2DX PrepFAST M5 autosampler (Elemental Scientific) equipped with 250 μL loop and using the 0.25 mL precision method provided by the manufacturer. Serumnorm™ (Sero, Norway) was used as a standard reference material and values for elements of interest were within 20% of the accepted value. Quantitative data analysis was conducted with Qtegra software and values were exported to Excel for further statistical analysis.

### RNAseq and Data Analysis

RNA preparation, library generation, and sequencing were done by BGI and are briefly described below. Total RNA concentration was determined with an Agilent 2100 Bio analyzer (Agilent RNA 6000 Nano Kit), as was QC metrics: RIN values, 28S/18S ratio and fragment length distribution.

Purification of the mRNA was done using a poly-T oligo immobilized to magnetic beads. Following purification, the mRNA is fragmented into pieces using divalent cations plus elevated temperature. The cleaved RNA fragments are copied into first strand cDNA using reverse transcriptase and random primers. Second strand cDNA synthesis was done using DNA Polymerase I and RNase H. These cDNA fragments then have the addition of a single ‘A’ base and subsequent ligation of adapter. The products are purified and enriched with PCR amplification. The PCR yield was quantified using a Qubit and samples were pooled together to make a single strand DNA circle (ssDNA circle), which produced the final library. DNA nanoballs (DNBs) were generated with the ssDNA circle by rolling circle replication (RCR) to enlarge the fluorescent signals at the sequencing process. The DNBs were loaded into the patterned nanoarrays and pairend reads of 100 bp were read through on the DNBseq platform. The DNBseq platform combines the DNA nanoball-based nanoarrays and stepwise sequencing using Combinational Probe-Anchor Synthesis Sequencing Method. On average, we generated about 4.41 Gb bases per sample. The average mapping ratio with reference genome was 92.56%, the average mapping ratio with gene was 72.22%; 17,240 genes were identified. 16,875 novel transcripts were identified. Read quality metrics: 4.56% of the total amount of reads contained more than 5% unknown N base; 5.92% of the total amount of reads which contained adaptors; 1.38% of the total reads were considered low quality meaning more than 20% of bases in the total read have quality score lower than 15. 88.14% of total amount of reads were considered clean reads and used for further analysis.

Analysis of sequencing reads. The sequencing reads were uploaded to the Galaxy web platform, and we used the public server at usegalaxy.eu to analyze the data (72). FastQC was performed to remove samples of poor quality (73). All mapping was performed using Galaxy server (v. 19.09) running Hisat2 (Galaxy Version 2.1.0+galaxy5), FeatureCounts (Galaxy Version 1.6.2), and DeSeq2 (Galaxy Version 2.11.40.2) (74–76). The Genome Reference Consortium build of the reference sequence (GRCh38/hg38) and the GTF files (NCBI) were used and can be acquired from iGenome (Illumina). Hisat2 was run with the following parameters: paired-end, unstranded, default settings were used except for a GTF file was used for transcript assembly. Alignments were visualized using IGV viewer (IGV-Web app version 1.7.0, igv.js version 2.10.5) with Ensembl v90 annotation file and Human (GRCh38/hg38) genome (77, 78).

The aligned SAM/BAM files were processed using Featurecounts (Default settings except used GRCh38 GTF file and output for DESeq2 and gene length file). FeatureCounts output files and raw read files are publicly available (GEO with accession GSE192392). The FeatureCounts compiled file is GSE192392_RawCountsCompiled. Gene counts were normalized using DESeq2 (Love et al., 2014) followed by a regularized log transformation. Differential Expression was determined using DESeq2 with the following settings: Factors were cell type, pairwise comparisons between mutant cell lines versus control line was done, output all normalized tables, size estimation was the standard median ratio, fit type was parametric, outliers were filtered using a Cook’s distance cutoff.

### Single Cell RNAseq

Single cell RNA seq data described by Yao et al were downloaded from [the Allen Institute Portal](https://portal.brain-map.org/atlases-and-data/rnaseq) and processed as previously described by us to extract information about SLC30A10 (79, 80).

### Synergy Analysis

Cells were counted using and automated Bio-Rad cell counter (Bio-Rad, TC20, 1450102). 2000 cells per well were plated on 96 well assay plates (Costar, 3603). Some wells were left intentionally blank and used as negative controls. The plates were then placed in the incubator at 37°C and 10% CO_2_. The following day the media on the cells were replaced with media containing a combination of manganese chloride (MP Biomedicals, 155334) and Actinonin (Sigma, A6671), Rotenone (Sigma, R8875), IMT1B (MedChemExpress, LDC203974) or Doxycycline (Sigma, D9891). The plates were placed back in the incubator overnight. On the day of the measurements media from the plate was aspirated and Resazurin reagent (R&D Systems, AR002) diluted 1:10 in media was added to each well. The plates were placed back in the incubator for 2 hours. Fluorescence was measured using a microplate reader (BioTek, Synergy HT; excitation at 530-570 nm and emission maximum at 580-590 nm) and the data was imported and analyzed using the BioTek Gen5 3.11 software. Synergy calculations were performed using the ZIP score with the SynergyFinder engine https://synergyfinder.org/ (48, 49).

### Blue Native Electrophoresis

Starting material was two 150mm dishes with cells at 80-90% confluency for each condition. The cells were released with trypsin and the pellet washed with PBS. Crude mitochondria were enriched according to Wieckowski et al.(81). Briefly, cells were homogenized in isolation buffer (225-mM mannitol, 75-mM sucrose, 0.1-mM EGTA and 30-mM Tris-HCl pH 7.4) with 20 strokes in a Potter-Elvehjem homogenizer at 6,000rpm, 4ºC. Unbroken cells and nuclei were collected by centrifugation at 600g for 5min and mitochondria recovered from this supernatant by centrifugation at 7,000g for 10 min. After 1 wash of this pellet, membranes were solubilized in 1.5 M aminocaproic acid, 50 mM Bis-Tris pH 7.0 buffer with antiproteases and 4 g/g digitonin. Proteins were separated by blue native electrophoresis in 3-12% gradient gels (82, 83) using 10mg/ml Ferritin (404 and 880 kDa, Sigma F4503) and BSA (66 and 132kDa) as molecular weight standards. Some samples were further separated in a second dimension 10% denaturing SDS-PAGE gels.

### Cell survival

On day zero, cells were plated at a density of 2,000 cells/well for HAP1 and 7,500 for SH-SY5Y. At day 1 cells were incubated with the indicated concentration of manganese and/or inhibitors. After 3 days of treatment, survival was measured following a 2h incubation with Alamar Blue or fixed with 100% methanol and then stained with 0.1% Crystal Violet. Percentage survival was calculated relative to untreated cells.

### Drosophila survival

Drosophila stocks obtained from the obtained from the Bloomington Drosophila Stock Center are as follows:

For over-expression of *scully:* P{UAS-scu.Exel}1, y1 w1118
*scully* RNAi: y1 v1; P{TRiP.HMS02305}attP40
background control: y[1] v[1]; P{y[+t7.7]=CaryP}attP40
actin GAL4 driver: y[1] w[*]; P{w[+mC]=Act5C-GAL4}17bFO1/TM6B, Tb[1]

Flies were reared on standard molasses food (Genesse Scientific) at 25°C in a 12hr light: 12hr dark cycle. Male and females were collected and aged 1 week. 7-10 male or female flies per replicate were transferred to empty vials and starved for 3 hours. Flies were then transferred to empty vials with a filter disc soaked in 200 μl of a 5% glucose, 10mM MnCl_2_ tetrahydrate solution. The number of dead flies was assessed every 24hrs.

### Promoter methylation analysis

DNA methylation was quantitatively assessed with the EpitTYPER (39) by CD-Genomics (New Jersey, USA). Potential CpG islands were identified in the promoter region 206764655 to 206767200 DMR (GRCh38.p13) of FASTKD2 gene obtained from the UCSC genome browser (http://genome.ucsc.edu/). Genomic DNA isolated from HAP1 SLC30A10 wild type and knockout cells was treated with bisulfite and then PCR amplified employing primers spanning 3 regions of interest:

5’ primer sequence: aggaagagag TGTTTTGTGTTTTATGAGATTGAAA
3’ primer sequence: cagtaatacgactcactatagggagaaggct CCAAAAAAAATAACTAATTTCCCTACA
5’ primer sequence: aggaagagag TATATGTTTTTTGATTTTTGAGGGAA
3’ primer sequence: cagtaatacgactcactatagggagaaggct AAAAAATCTTATACAACCAACCAACC
5’ primer sequence: aggaagagag GAAAATGGGGAATTTATGGAGTTAT
3’ primer sequence: cagtaatacgactcactatagggagaaggct CCTAAAATCAAAAATTCAAAACCAA

The amplicons were transcribed into RNA and fragmented by RNAseA cleavage. Mass of the fragments was estimated by matrix Assisted Laser Desorption-Ionization Time of Flight. The DNA methylation percentage of a given CpG is calculated by dividing the surface area of the peak representing the methylated fragment by the total surface area of the peaks of both the methylated and unmethylated fragment.

### Mitochondrial RNA transcript junction measurements

RNA junctions were profiled employing transcript counting on a Nanostring nCounter Platform with a set of custom designed probes (34) for multiples detection of all junctions of human polycistronic mitochondrial transcripts, some non-coding RNA regions, and genes to standardize the samples. Clathrin was the gene chosen for standardization across samples. Cells were plated in 10cm dishes and collected with 1ml Trizol when they reached 80% confluency. RNA was purified and quality was controlled employing an Agilent Bioanalyzer before hybridization and counting.

### Oxygen consumption

Extracellular flux analysis on the Seahorse XFe96 Analyzer (Seahorse Bioscience) following manufacturer recommendations. XFe96 extracellular flux assay kit probes (Seahorse Bioscience 102601-100) were incubated with the included calibration solution overnight at 37°C under non-CO_2_-injected conditions. Cells were trypsinized, counted (Bio-Rad TC20 automated Cell Counter), and seeded into XFe96 cell culture microplates (Seahorse Bioscience 101085-004) at 32,000 cells/well for SHSY5Y and 40,000 for HAP1 cells and incubated at 37°C with 10% CO_2_ in complete culture media for 24 h before initialization of the mitochondrial stress test. The next day, wells were washed twice in Seahorse stress test media consisting of Seahorse XF base media (Seahorse Bioscience 102353-100), 2 mM L-glutamine (HyClone SH30034.01), 1 mM sodium pyruvate (Sigma S8636), and 10 mM D-glucose (Sigma G8769), pH7.4 before final volume added and then incubated at 37°C in non-CO_2_-injected conditions for 1 h before stress test. Seahorse injection ports were filled with 10-fold concentrated solution of oligomycin A (Sigma 75 351), carbonyl cyanide-4-(trifluoromethoxy)phenylhydrazone (FCCP; Sigma C2920), and rotenone (Sigma R8875)/antimycin A (Sigma A8674) for final testing conditions of oligomycin (1.0 μM), FCCP (0.125 μM for HAP1 0.25uM for SH-SY5Y), rotenone (0.5 μM), and antimycin A (0.5 μM). The flux analyzer protocol conditions consisted of three basal read cycles, and three reads following each injection of oligomycin A, FCCP, and rotenone plus antimycin A. Each read cycle consisted of a 3-min mix cycle, followed by a 3-min read cycle where oxygen and pH levels were determined over time. The Seahorse Wave Software version 2.2.0.276 was used for data analysis of oxygen consumption rates. Experiments were repeated in triplicate. Non-mitochondrial respiration was determined as the lowest oxygen consumption rate following injection of rotenone plus antimycin A. Basal respiration was calculated from the oxygen consumption rate just before oligomycin injection minus the non-mitochondrial respiration. Oligomycin A sensitivity was calculated as the difference in oxygen consumption rates just before oligomycin injection to the minimum oxygen consumption rate following oligomycin injection but before FCCP injection.

### Bioinformatic Analyses and Statistical Analyses

Gene ontology analyses were performed with Cluego. ClueGo v2.58 run on Cytoscape v3.8.2 (84, 85). ClueGo was run querying GO BP and KEGG considering all evidence, Medium Level of Network Specificity, and selecting pathways with a Bonferroni corrected p value <0.05. ClueGo was run with Go Term Fusion. PCI and 2D-tSNE were calculated using Qlucore Omics Explorer Version 3.6(33) normalizing log_2_ data to a mean of 0 and a variance of 1. 2D-tSNE was calculated with a perplexity of 5. All other statistical analyses were performed with Prism v9.2.0(283) using two tailed statistics and Alpha of 0.05.

## Data Availability

The mass spectrometry proteomics data have been deposited to the ProteomeXchange Consortium via the PRIDE (86) partner repository with dataset identifier: PXD030502 Project DOI: 10.6019/PXD030502

RNAseq data were deposited in GEO with accession GSE192392

## Acknowledgements

This work was supported by grants from the National Institutes of Health 1RF1AG060285 to VF, 1P50NS098685 Udall Parkinson Center to AG. Fly stocks obtained from the Bloomington *Drosophila* Stock Center (NIH P40OD018537) were used in this study. This study was supported in part by the Emory Integrated Genomics Core (EIGC), which is subsidized by the Emory University School of Medicine. VF is grateful for mitochondria provided by Maria Olga Gonzalez

## Supplementary Figure Legends

**Fig. S1.**
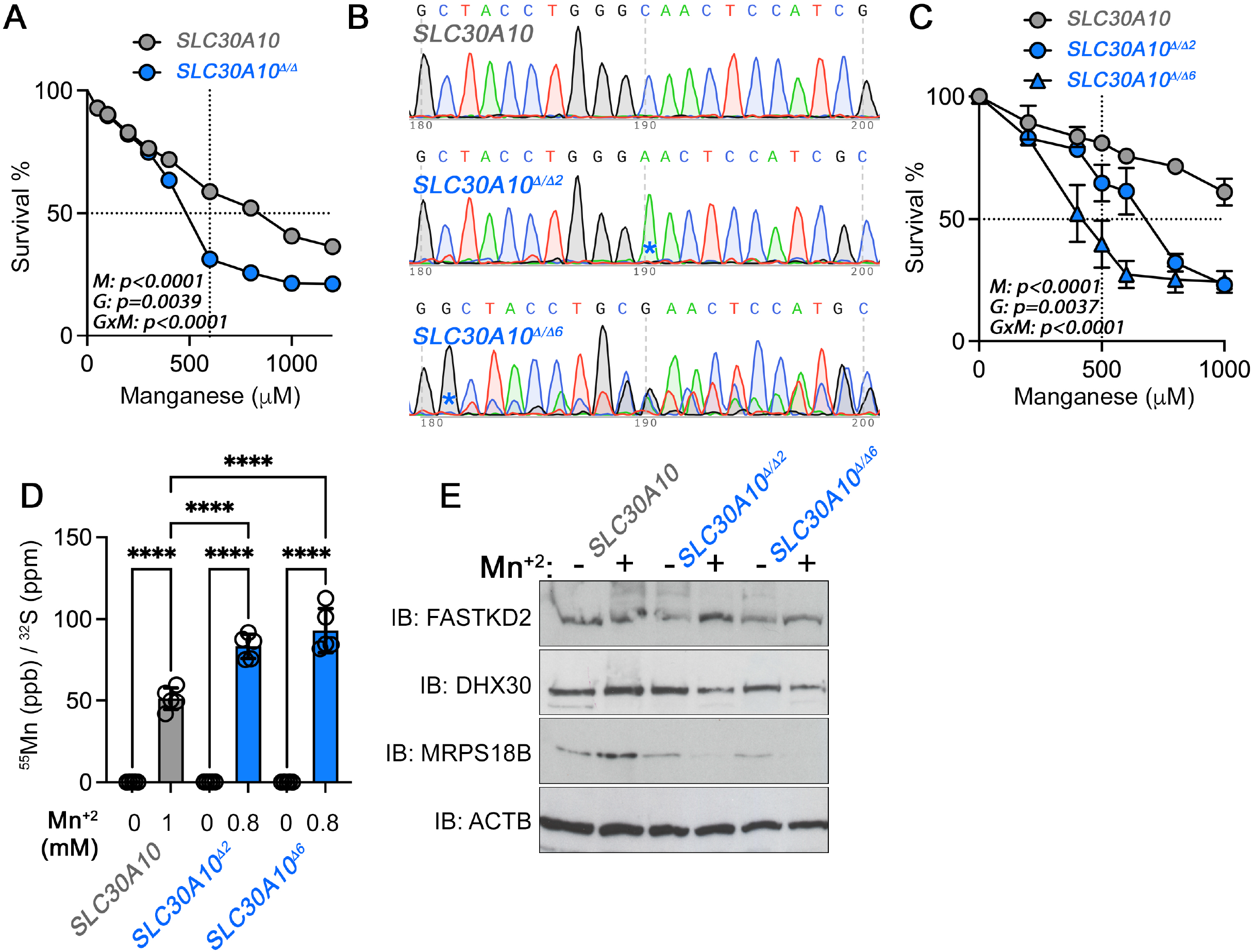
Generation of SLC30A10 Mutant Neuroblastoma Cells. **A**. Cell survival analysis of SLC30A10 mutants with increasing concentrations of manganese. Mutant cells correspond to a 95% CRISPR knock out SH-5YSY cell pool. Average ± SEM, n=3. Two-Way ANOVA followed by Šydák’s multiple corrections. Error bars are smaller than the symbol. **B**. Sequence chromatograms of wild type and two SH-5YSY CRISPR clones isolated from the pool in panel B. Asteriks mark the beginning of the mutant sequence. **C**. Cell survival analysis of SLC30A10 mutant clones shown in panel B and their controls. Average ± SEM, n=3. Two-Way ANOVA followed by Šydák’s multiple corrections. **D**. ICP mass spectrometry determination of manganese in wild type and clones presented in panels B-C. Vehicle or manganese was added overnight. Data were normalized to total sulfur content. Average ± SEM, n=5, Two-Way ANOVA followed by Bonferroni corrections. **E**. Immunoblot analysis of SLC30A10 mutant clones blotted for granule components. Actin (ACTB) was used as a loading control.

**Fig. S2.**
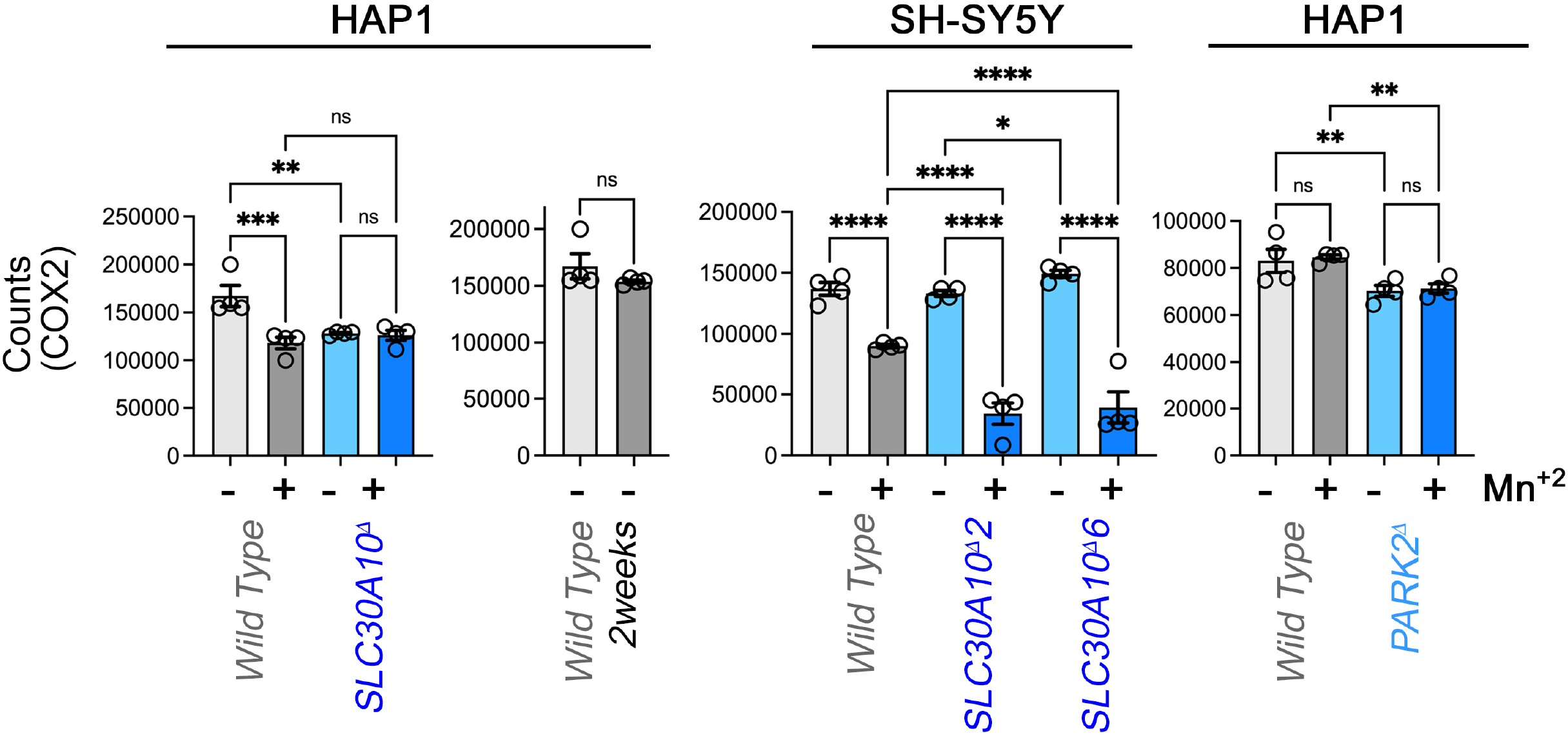
Expression of the Mitochondrial Encoded COX2 Subunit. Nanostring analysis of the COX2 mitochondrial mRNA using the Mitostring panel, see figure legend 4 for details. Wild type cells and mutants were incubated either in the presence or in the absence of manganese overnight. Average ± SEM, n=4. One-Way ANOVA followed by Benjamini, Krieger and Yekutieli corrections.

**Figure S3.**
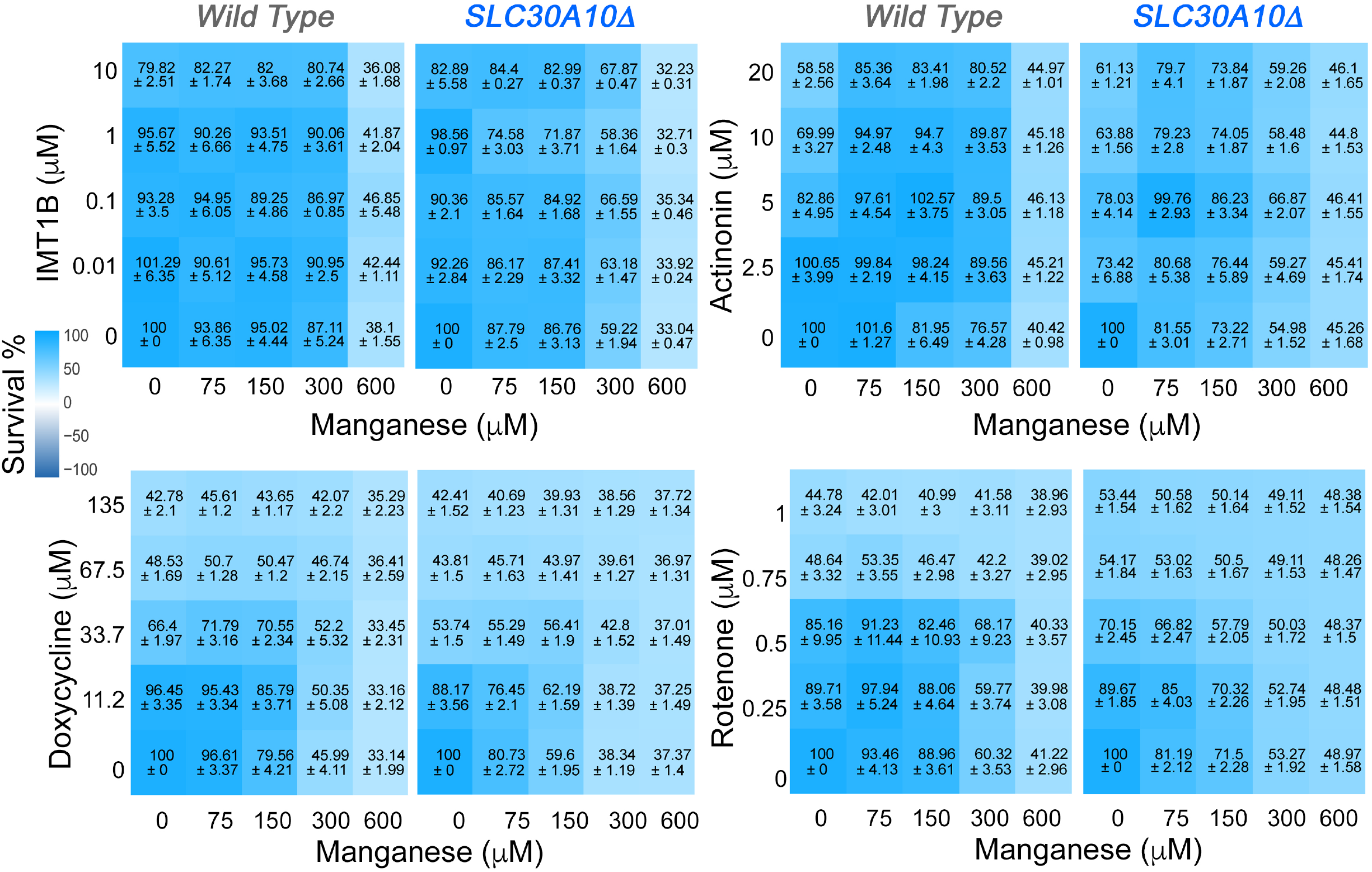
Dose Response Maps for Manganese-Drug interaction in Wild Type and SLC30A10 Mutant Cells. Heat Map represents cell survival after manganese-drug pair additions. Number in each cell represent average ± SEM. These values were used to generate the Synergy maps in Figure 6F.

**Figure S4.**
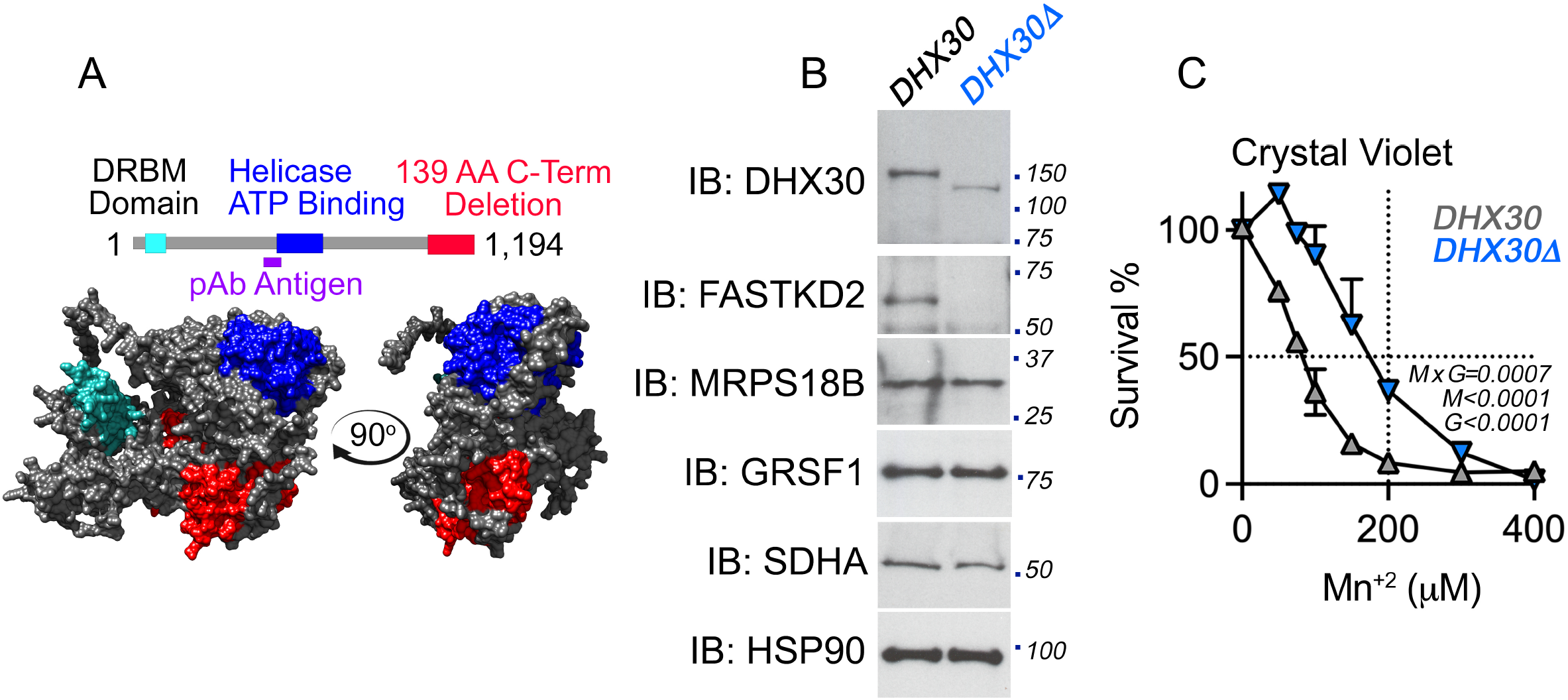
Characterization of a DHX30 Hypomorph Mutant HAP1 Cell Line. HAP1 cells were edited by CRISPR-Cas9 targeting exon 7 of transcript NM_138615. Mutation creates a 26 bp deletion in exon 7. **A**. shows diagram of DHX30 with DRBM, helicase, and predicted deletion color coded teal, blue and red; respectively. AlphaFold presumptive model of human DHX30 with domains marked in the same colors as in A (https://alphafold.ebi.ac.uk/entry/Q7L2E3). The antigen used to raise a DHX30 antibody (residues 350-400) is mark in purple. Note that the red deleted segment sits on a predicted chief globular domain of DHX30. B. Immunoblots of wild type and DHX30 mutant cells show decreased levels of a truncated DHX30 protein and undetectable FASTKD2 expression. **C.** Cell survival analysis of wild type and DHX30 mutant cells exposed to increasing concentrations of manganese. Assay was performed with Crystal Violet. Average ± SEM, n=2. Two-Way ANOVA followed by Bonferroni corrections.

**Fig. S5.**
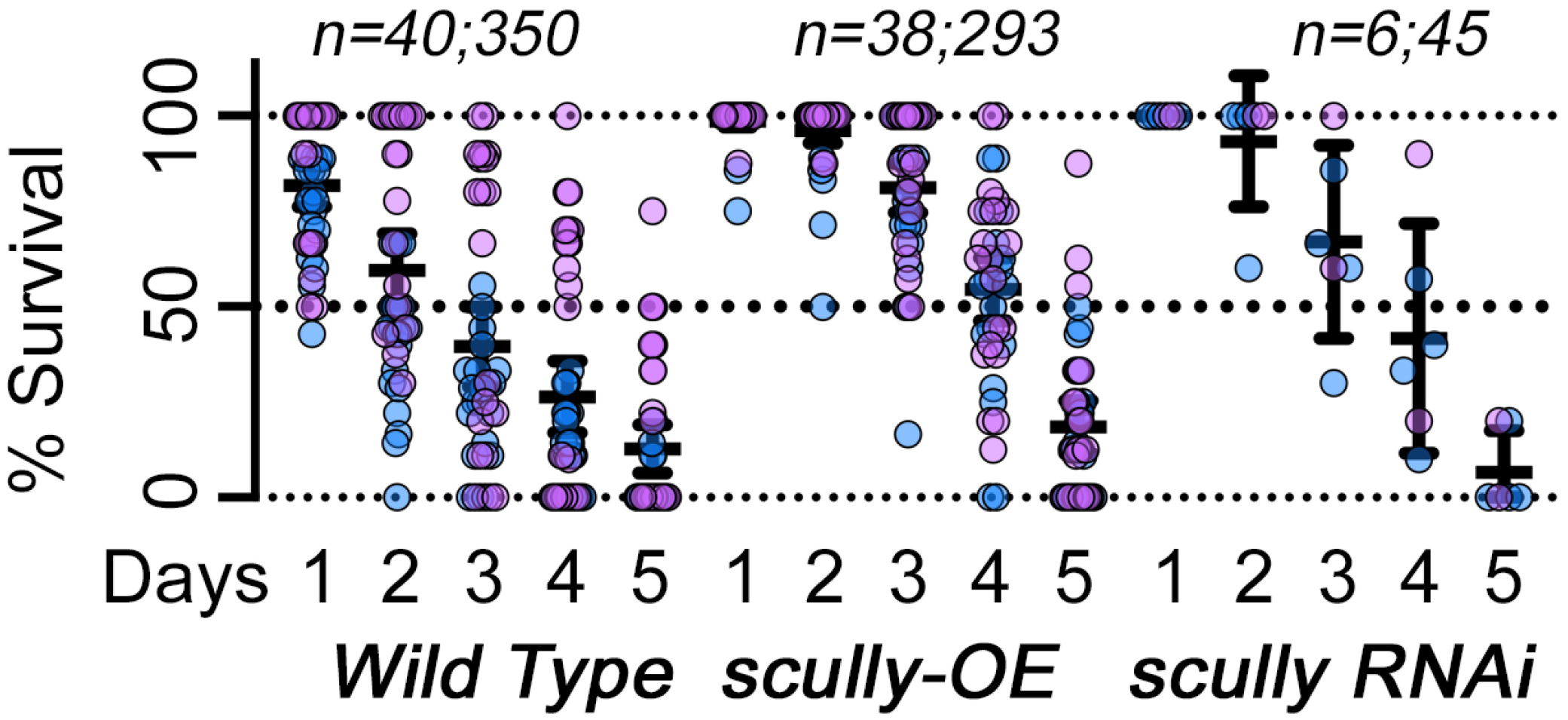
Survival of Adult Drosophila Exposed to Dietary Manganese is Increased after Disruption of the Mitochondrial RNA Granule. Wild type, *scully* RNAi, or *scully* overexpressing animals were fed a manganese containing diet for 5 days. Average ± 95% CI, blue symbols depict males and purple symbols depict female flies. There was not difference between sexes. n=number of independent experiments per genotype, number of total animals studied per genotype. Number of replicates were as follows: male controls n=20, male *scully* RNAi n=4, male *scully* over-expression n=18, female controls n=20, female *scully* RNAi n=2, female *scully* over-expression n=20.

**Supplementary Table 1. TMT Mass Spectrometry Results.**

**Supplementary Table 2. Integrated ClueGo Ontology Results**

**Supplementary Table 2. DeSeq2 RNAseq Results**

